# A novel motif in calcimembrin/C16orf74 dictates multimeric dephosphorylation by calcineurin

**DOI:** 10.1101/2024.05.12.593783

**Authors:** Devin A. Bradburn, Joana C. Reis, R.Yvette Moreno, Shariq Qayyum, Thibault Viennet, Haribabu Arthanari, Martha S. Cyert

**Affiliations:** Department of Biology, Stanford University, Stanford, CA 94305, USA; Department of Cancer Biology, Dana-Farber Cancer Institute, Boston, Massachusetts, USA; Department of Biological Chemistry and Molecular Pharmacology, Harvard Medical School, Boston, Massachusetts, USA; Men’s Health: Aging & Metabolism, Brigham and Women’s Hospital, Harvard Medical School, Boston, MA, USA; Cardiovascular Research Center, Department of Medicine, Massachusetts General Hospital and Harvard Medical School, Boston, MA, USA; Department of Chemistry and iNANO, Aarhus University, Aarhus, Denmark

## Abstract

Calcineurin, the Ca^2+^/calmodulin-activated protein phosphatase, recognizes substrates and regulators via short linear motifs, PxIxIT and LxVP, which dock to distinct sites on calcineurin to determine enzyme distribution and catalysis, respectively. Calcimembrin/C16orf74 (CLMB), an intrinsically disordered microprotein whose expression correlates with poor cancer outcomes, targets calcineurin to membranes where it may promote oncogenesis by shaping calcineurin signaling. We show that CLMB associates with membranes via lipidation, i.e. N-myristoylation and reversible S-acylation. Furthermore, CLMB contains an unusual composite ‘LxVPxIxIT’ motif, that binds the PxIxIT-docking site on calcineurin with extraordinarily high affinity when phosphorylated, ^33^LDVPDIIITPP(p)T^44^. Calcineurin dephosphorylates CLMB to decrease this affinity, but Thr44 is protected from dephosphorylation when PxIxIT-bound. We propose that CLMB is dephosphorylated in multimeric complexes, where one PxIxIT-bound CLMB recruits calcineurin to membranes, allowing a second CLMB to engage via its LxVP motif to be dephosphorylated. *In vivo* and *in vitro* data, including nuclear magnetic resonance (NMR) analyses of CLMB-calcineurin complexes, support this model. Thus, CLMB with its composite motif imposes unique properties to calcineurin signaling at membranes including sensitivity to CLMB:calcineurin ratios, CLMB phosphorylation and dynamic S-acylation.

## Introduction

Calcium (Ca^2+^) ions are spatially and temporally controlled in cells to regulate a wide array of biological processes ^1^. Ca^2+^ signaling frequently occurs within membrane-associated microdomains that contain both effector proteins and their substrates. Thus, subcellular localization of calcineurin, the only Ca^2+^-calmodulin (CaM) regulated serine/threonine protein phosphatase, dictates its signaling specificity^2^. Calcineurin (CNA/B) is an obligate heterodimer composed of catalytic (CNA) and regulatory (CNB) subunits that is activated when Ca^2+^/calmodulin binding relieves autoinhibition mediated by the CNA C-terminal domain^3^. Calcineurin interacts with partners via two short linear motifs (SLiMs), LxVP and PxIxIT, which cooperate to promote dephosphorylation. PxIxIT binds to CNA independent of Ca^2+^and anchors calcineurin to substrates and regulators. In contrast, LxVP binds to a site composed of residues in both CNA and CNB and orients substrates for dephosphorylation at a separate catalytic site, marked by Zn^2+^ and Fe^2+^ ions^2, 3^. LxVP binding occurs only after activation by Ca^2+^/CaM and is blocked by autoinhibitory sequences in CNA as well as the immunosuppressant drugs, Cyclosporin A and FK506^4, 5^.

Systematic discovery of PxIxIT and LxVP SLiMs has significantly expanded our knowledge of calcineurin signaling^6, 7, 8, 9^ and surprisingly identified at least 31 human proteins that harbor composite calcineurin-binding motifs containing contiguous LxVP and PxIxIT sequences that share a proline (LxVPxIxIT)^9^. Although each SLiM in this composite motif can bind to its cognate site, they are unlikely to do so simultaneously, as their docking surfaces are ∼60 Å apart on calcineurin^4, 9^. Thus, how or if these motifs promote dephosphorylation is unclear.

Here we elucidate how C16orf74, a short unstructured microprotein associated with poor outcomes in melanoma, pancreatic and head and neck cancer, is regulated by calcineurin via a composite LxVPxIxIT motif ^9, 10^. We name C16orf74 calcimembrin (CLMB), because it associates with membranes via lipidation (N-myristoylation and S-acylation) and recruits calcineurin to these sites. CLMB is phosphorylated at Thr 44 (pThr44) and binds calcineurin with exceptionally high affinity because the flanking LxVP and pThr44 features (LxVPxIxITxx(p)T) enhance PxIxT-mediated binding to CNA. Calcineurin also dephosphorylates pThr44, but this residue is protected when CLMB is bound to the PxIxT-docking surface. Thus, we propose that dephosphorylation occurs in multimers, with one CNA/B bound to two CLMB molecules at the PxIxIT and LxVP-docking sites, respectively. NMR confirms formation of these complexes *in vitro*. Furthermore, this multimeric model predicts that CLMB dephosphorylation varies depending on the amount of CLMB relative to calcineurin, which we demonstrate via in cell analyses and kinetic modeling of dephosphorylation *in vitro*. In summary, this work reveals how CLMB can dynamically shape calcineurin signaling through a mechanism dictated by its composite motif and reversible association with membranes.

## Results

### Calcimembrin is myristoylated and S-acylated

CLMB lacks predicted transmembrane domains but may associate with membranes via lipidation (Fig. 1a)^10^. CLMB contains a glycine at position 2 (Gly2), a site frequently modified by myristate, a 14-carbon lipid whose co-translational addition can promote protein association with membranes^11^. CLMB also has two cysteines near in its N-terminus, Cys7 and Cys14, one of which, Cys7, is predicted to be modified by S-acylation, the reversible addition of a fatty acid, usually a 16-carbon palmitate, that regulates membrane localization^12^. To determine if CLMB is lipidated, we metabolically labeled HEK293 Flp-In T-REx cells inducibly expressing wildtype CLMB-FLAG (CLMB_WT_) with YnMyr and 17-ODYA, alkyne-containing analogs of myristate and palmitate respectively^13^. Following immunoprecipitation, analog incorporation was detected via addition of biotin using CLICK-chemistry followed by visualization with streptavidin (Fig. 1b). CLMB_WT_ incorporated YnMyr and 17-ODYA, indicating that the protein is both myristoylated and palmitoylated. To identify which residues were lipidated, we mutated proposed sites of S-acylation (CLMB_C7,14S_), and myristoylation (CLMB_G2A_ (Fig. 1b). The mutants were expressed at low levels suggesting instability, which we countered by including a proteasome inhibitor (MG132) in the analyses. CLMB_C7,14S_ showed no 17-ODYA incorporation but maintained YnMyr suggesting that Cys7 and/or Cys14 are S-acylated. By contrast, neither lipid modification was detected for CLMB_G2A_, which confirms that Gly2 is myristoylated. Because myristoylation can promote S-acylation by anchoring a protein to the membrane^14^, we then examined CLMB_G2A_ for S-acylation with increased sensitivity by including palmostatin B (PalmB), a pan-inhibitor of thioesterases that remove S-acylation^15^. In the presence of this inhibitor, 17-ODYA incorporation into both CLMB_WT_ and CLMB_G2A_ was observed (Supplementary Fig.1), indicating that CLMB_G2A_ is S-acylated, but with reduced efficiency.

**Fig. 1.**
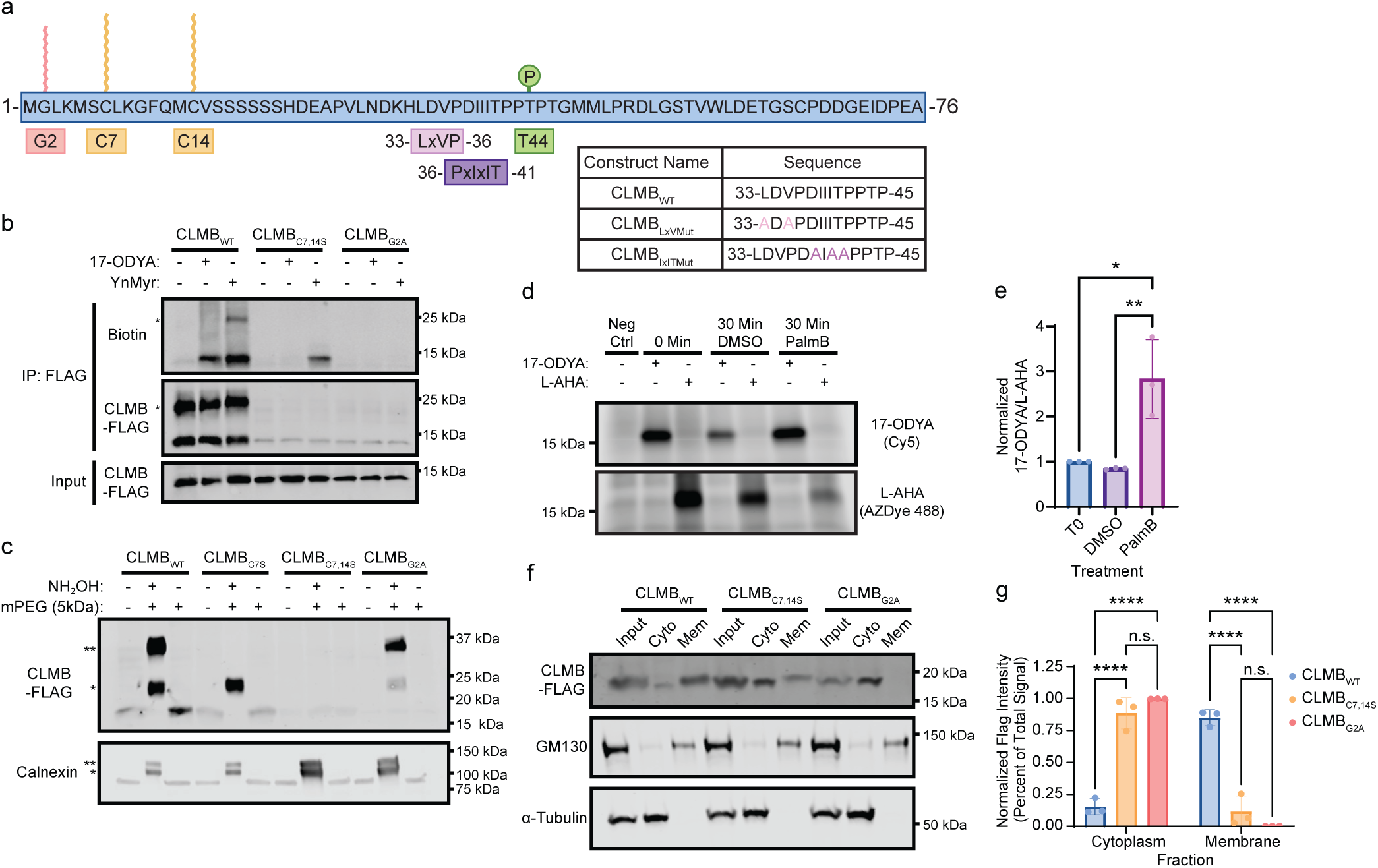
Calcimembrin localizes to membranes via protein lipidation. **a.** CLMB schematic showing N-myristoylation (red); S-acylation (yellow), phosphorylation (green), calcineurin-binding motifs LxVP (pink) and PxIxIT (purple). Sequences of wild-type and mutated regions shown. **b.** Representative immunoblot showing metabolic labeling of CLMB-Flag (wild-type or indicated mutants) with palmitate (17-ODYA) or myristate (YnMyr) analogs, analyzed via CLICK chemistry with biotin-azide and detection with IRdye 800CW streptavidin (“Biotin”). Inputs and immunoprecipitated (IP) samples are shown. Asterisks indicate CLMB dimers. (n = 3 independent experiments). **c.** Representative immunoblot showing S-acylation of CLMB-FLAG (wild-type or indicated mutants) and endogenous calnexin (positive control) detected via Acyl-PEG exchange. Asterisks show PEGylation events. Input, cleaved (NH_2_OH) and PEGylated (mPEG), and non-cleaved PEGylated conditions are indicated (n = 3 independent experiments). **d.** Representative in-gel fluorescence scan showing pulse-chase analysis of CLMB-FLAG using palmitate (17-ODYA) or methionine (L-AHA) analogs detected via CLICK chemistry with Cy5 or AZDye 488, respectively. Incorporation into CLMB-FLAG with no analog (Neg Ctrl), at 0 or 30 minutes after unlabeled chase with DMSO (control) or PalmB. shown (n = 3 independent experiments). **e.** Data from (**d**) showing mean ± SEM of 17-ODYA signal normalized to L-AHA; * p < 0.05, T0 vs PalmB p = 0.0103, ** p < 0.01, DMSO vs PalmB p = 0.007**. f**. Representative immunoblot showing sub-cellular fractionation of CLMB-FLAG (wild-type or indicated mutants) with input, cytoplasmic (cyto), and membrane (mem) fractions. GM130 and α-tubulin mark mem and cyto fractions, respectively (n= 3 independent experiments). **g**. Data from (**f**) showing mean ± SEM cyto or mem signals normalized to total (cyto + mem); n.s. not significant, **** p < .0001. CLMBC_7,14S_ vs CLMB_G2A_: cytoplasm, p = 0.238, membrane p = 0.238. For (**e, g**) p-values were calculated by two-way ANOVA corrected by Tukey’s multiple comparison.

To determine if CLMB-FLAG is dually S-acylated at Cys7 and Cys14, we used acyl-PEG exchange (APE) in which S-acylated groups are cleaved using hydroxylamine (NH_2_OH) and exchanged for a PEGylated molecule that increases protein mass by 5 kDa for each S-acylated residue^16^ (Fig.1c). Endogenous calnexin, a dually acylated protein, served as a positive control for each sample. For CLMB_WT_, two slower migrating bands were observed. CLMB_C7S_ lacked the upper band, while CLMB_C7,14S_ was unmodified, indicating that Cys7 and Cys14 are S-acylated in CLMB_WT_ (Fig. 1c). CLMB_G2A_ displayed two sites of S-acylation, but with reduced intensity relative to CLMB_WT_, consistent with results from metabolic labeling (Fig. 1b). Together these findings establish that CLMB is myristoylated at Gly2 and S-acylated at Cys7 and Cys14.

### Dynamic S-acylation of CLMB mediates membrane association

To examine whether S-acylation of CLMB is dynamic^17^, we carried out a pair of pulse-chase analyses with two metabolic labels (Fig. 1d, e). L-AHA, an azide-containing methionine analog detected CLMB-FLAG protein levels, while in parallel, 17-ODYA detected S-acylation. CLMB-FLAG was immunopurified (IP) after labeling for one hour followed by a 30-minute chase with DMSO or PalmB. Quantifying the 17-ODYA to L-AHA ratio revealed that palmitate incorporation into CLMB increased ∼3-fold when thioesterases were inhibited (Fig, 1e, PalmB vs DMSO). This indicates that S-acylation of CLMB turns over rapidly relative to its rate of degradation.

To determine if lipidation mediates CLMB membrane recruitment, we analyzed the localization of CLMB_WT_ and lipidation defective mutants, CLMB_G2A_ and CLMB_C7,14S_. Detergent-assisted subcellular fractionation showed that CLMB_WT_ was primarily membrane associated, appearing in fractions marked by GM130, a Golgi protein. By contrast, both mutants were predominantly cytosolic, appearing in fractions marked by α-tubulin (Fig. 1 f, g). Together these analyses indicate that N-myristoylation, in combination with dynamic dual S-acylation, promote the stable association of CLMB with membranes.

### The composite SLiM in CLMB mediates binding to and dephosphorylation by calcineurin

Next, we examined how each part of CLMB’s composite motif, LDVPDIIIT, contributes to calcineurin binding by generating mutants, CLMB_IxITMut_ (LDVPD**A**I**AA**) and CLMB_LxVMut_, (**A**D**A**PDIIIT) (Fig. 1a) that disrupt each motif independently without altering the shared proline. To examine binding of these mutants to calcineurin, the association of CLMB-FLAG with GFP-tagged calcineurin (CNA-GFP complexed with endogenous CNB) was assessed following purification of calcineurin from lysates of HEK293 cells co-transfected with these constructs (Fig. 2a, b). CLMB_WT_ but not CLMB_IxITMut_ co-purified with calcineurin, showing that this interaction was PxIxIT-dependent. Less calcineurin co-purified with CLMB_LxVMut_ vs. CLMB_WT_, showing that the LxVP also makes a modest contribution to calcineurin binding. Furthermore, when expressed and analyzed by SDS-polyacrylamide gel electrophoresis (SDS-PAGE), both mutants displayed two electrophoretic forms, with a slower migrating band also faintly observed for CLMB_WT_ (Fig. 2a, input). Phosphorylation, which often reduces electrophoretic mobility, has been reported for CLMB at Thr44 (pThr44) and less frequently at Thr46^18^. We confirmed that the slower migrating forms of CLMB contained pThr44, as they were absent in mutants lacking this residue (CLMB_T44A_, CLMB_T44A+LxVMut_, CLMB_T44A,T46A_), but not in CLMB_T46A_ (Fig. 2a, Supplemental Fig. 2a). These observations indicate that calcineurin binds to CLMB and dephosphorylates pThr44, which is disrupted when either the LxVP or PxIxIT portion of the composite motif is compromised.

**Fig. 2.**
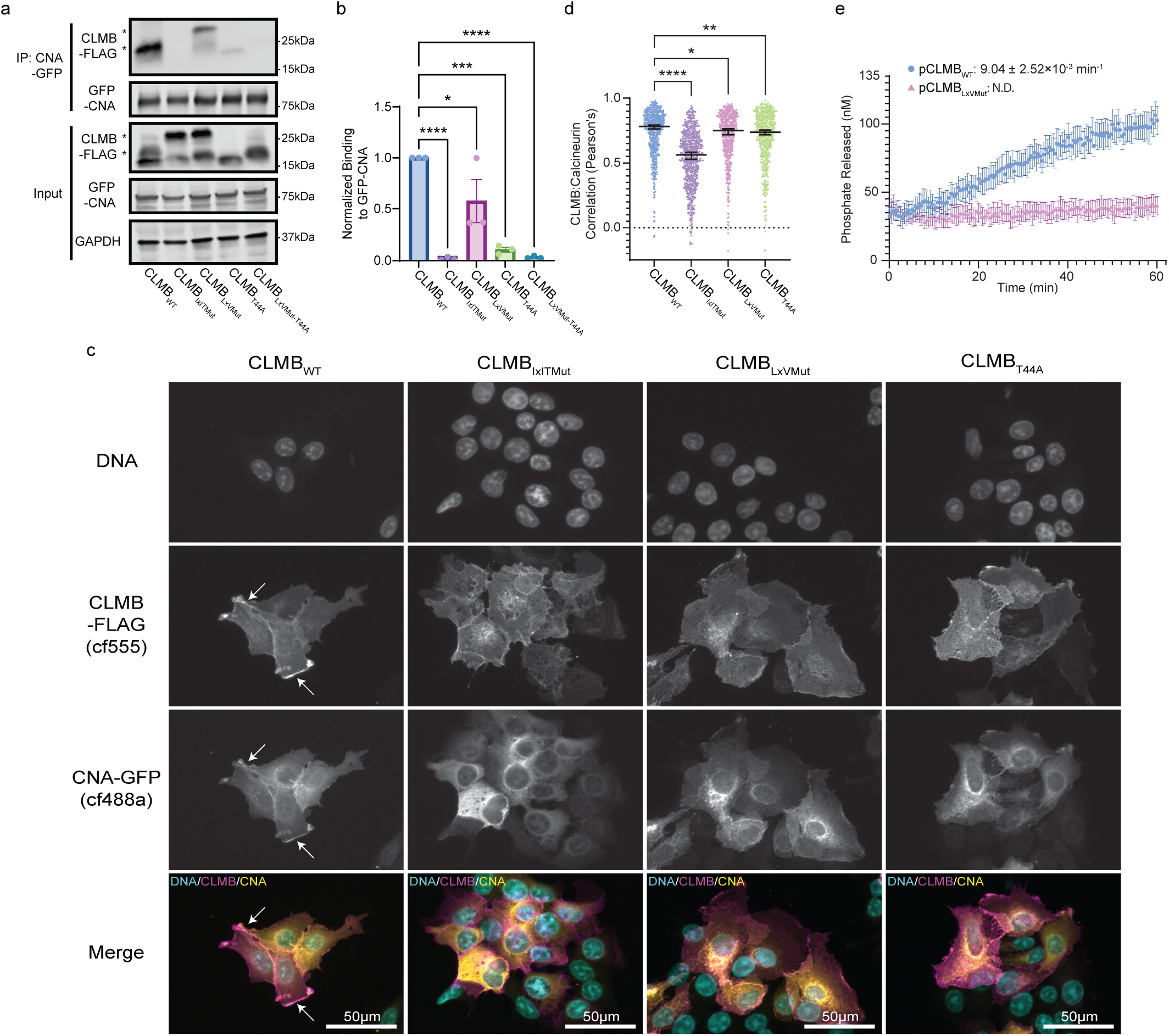
Composite LxVPxIxIT SLiM in calcimembrin mediates binding to and dephosphorylation by calcineurin. **a.** Representative immunoblot showing co-purification of CLMB-FLAG (wild-type and indicated mutants) with CNA-GFP. Inputs and purified (IP) samples shown (n = 3 independent experiments). Glyceraldehyde-3-Phosphate Dehydrogenase (GAPDH) was used as a loading control. **b.** Data from (**a**) showing mean ± SEM of co-purified CLMB-FLAG normalized to CLMB-FLAG input levels and precipitated CNA-GFP. Indicated p-values calculated by two-way ANOVA corrected by Dunnett’s multiple comparison test. **** p < 0.0001, *** p<0.001, * p<0.05; CLMB_WT_ vs CLMB_LxVmut_ p = 0.0325, CLMB_WT_ vs CLMB_T44A_ p = 0.0002. **c.** Representative images of MCF7 cells co-transfected with CLMB-FLAG (wild-type or indicated mutants) and CNA-GFP. Fixed cells immunostained with anti-FLAG (magenta), anti-GFP (yellow), and nuclei (DNA) with Hoechst 33342. Scale bar = 50 μm. Arrows indicate some examples of CLMB and CNA co-localization to plasma membrane. Data from (**c**) showing median with 95% CI Pearson’s correlation coefficient of CLMB-calcineurin colocalization. Each point represents a single cell (n ≥ 500 for each condition). p-values calculated by Kruskal-Wallis with Dunn’s multiple comparisons correction. * p < 0.05 (p = 0.0116), ** p < 0.01 (p = 0.0092), **** p < 0.0001. **e** Dephosphorylation of pCLMB_WT_ (blue) or pCLMB_LxVMut_ (pink) phosphopeptides with 20 nM CNA/B and 1 μM CaM. Mean ± SEM of phosphate released displayed for ≥ 4 replicates at each timepoint with reaction rate as shown.

Finally, we confirmed that pThr44 enhance binding to calcineurin, as the slower migrating (phosphorylated) forms of CLMB_WT_ and CLMB_LxVMut_ predominantly co-purified with calcineurin (Fig. 2a, IP)^10^. Indeed, CLMB_T44A_ displayed reduced binding to calcineurin, which was further reduced by mutating LxV (CLMB_T44A+LxVMut_) (Fig. 2a, b).

The contributions of PxIxIT, LxVP, and pThr44 to calcineurin binding were also examined by measuring the extent of CLMB co-localization with calcineurin (CNA-GFP complexed with endogenous CNB) via immunofluorescence (IF) (Fig. 2c, d). When co-transfected into MCF7 cells CLMB_WT_-FLAG and calcineurin colocalize at membranes (see regions marked by arrows in Fig. 2c). In contrast, when co-expressed with CLMB_IxITMut_, this colocalization was greatly reduced and calcineurin was predominantly cytosolic. CLMB_LxVMut_ and CLMB_T44A_ also displayed reduced colocalization with calcineurin (Fig. 2c, d), mirroring the trends observed in co-purification analyses (Fig. 2a, b). Thus, CLMB recruits calcineurin to membranes primarily via PxIxT-mediated binding.

Finally, dephosphorylation of pThr44 in CLMB by calcineurin was demonstrated *in vitro* using phosphorylated peptide substrates that encode residues 21-50 (Fig. 1a). The peptide from pCLMB_WT_ was robustly dephosphorylated compared to pCLMB_LxVMut_ (Fig. 2e) showing that, as for other calcineurin substrates, the LxVP motif promotes catalysis^19^. Finally, the presence of a functional LxVP motif in the CLMB_WT_ peptide was also confirmed by its ability to inhibit dephosphorylation of the RII peptide, an LxVP-containing calcineurin substrate derived from the regulatory subunit of protein kinase A (RII isoform)^19^ (Supplemental Fig. 2b). Together, these data show that both the LxVP and PxIxIT components of the composite motif are functional, that CLMB is a calcineurin substrate, and that pThr44 increases CLMB binding to calcineurin.

### Phosphorylated Thr44 and the LxVP motif enhance CLMB binding to the PxIxIT docking site

To quantitatively assess CLMB-calcineurin interaction, we used isothermal titration calorimetry (ITC) and fluorescence polarization anisotropy (FP) to measure binding of CLMB peptides containing the composite SLiM (residues 21-50) to purified, catalytically inactive CNA/B containing CNA_Trunc+H151Q_. This mutant CNA lacks the C-terminal autoinhibitory region, which allows LxVP motifs to bind to CNA/B in the absence of Ca^2+^/CaM^20^. ITC revealed that the phosphorylated CLMB_WT_ peptide (pCLMB_WT_) binds calcineurin at a one-to-one ratio with high affinity (*K*_D_= 37 nM), (Fig. 3a). FP using FITC-labelled peptides at pH 7.4 revealed that phosphorylation of Thr44 enhances CLMB binding to calcineurin ∼20-fold (*K*_D_ = 14.1 nM vs 291 nM) (Fig. 3b). Furthermore, the affinity of pCLMB_WT_ for calcineurin was similar over a pH range (6.9-8.0) which should change the protonation state of histidine residues (Supplemental Fig. 3a, b). This included the histidine immediately preceding the LxVP (H32) which is expected to engage with calcineurin during LxVP docking^7^. The concordance between measurements made with CLMB peptide that is tag-free (ITC) vs. tagged with fluorescein isothiocyanate (FITC) (FP) (37 vs 14.1 nM for pCLMB_WT_) (Fig. 3a, b), shows that the tag does not significantly affect binding affinity. Given the high-throughput capability of FP, this method was used for the remaining comparisons. To test if the high affinity pCLMB_WT_-calcineurin binding observed was PxIxIT-mediated, we used FP to assess the ability of CLMB peptides to compete with a well characterized PxIxIT peptide, FITC-PVIVIT ^21, 22^, for calcineurin binding (Fig. 3c, e). pCLMB_WT_ competed with PVIVIT most effectively, with peptides lacking the LxVP motif (pCLMB_LxVMut_), or pThr44 (CLMB_WT_) showing reduced competition (Fig. 3c, e). pCLMB_IxITMut_ lacking the PxIxIT motif was unable to displace PVIVIT from CNA/B. When these experiments were repeated using FITC-pCLMB_WT_ peptide in place of FITC-PVIVIT (Fig. 3d, e), a similar pattern was observed, although phosphorylation of T44 (pCLMB_WT_ vs CLMB_WT_) caused a much greater reduction in Ki (30X vs 3X for PVIVIT competition). Importantly, CLMB peptides accurately mimic full length CLMB binding to calcineurin as both caused equivalent displacement of FITC-pCLMB_WT_ (Supplemental Fig. 3c,d). Altogether, these data show that, like PVIVIT, pCLMB_WT_ peptides bind to the PxIxIT docking groove on CNA and that flanking residues, i.e. LxV and pThr44, enhance binding to form an extended LxVPxIxITxxT(p) sequence with exceptionally high affinity for calcineurin.

**Fig. 3.**
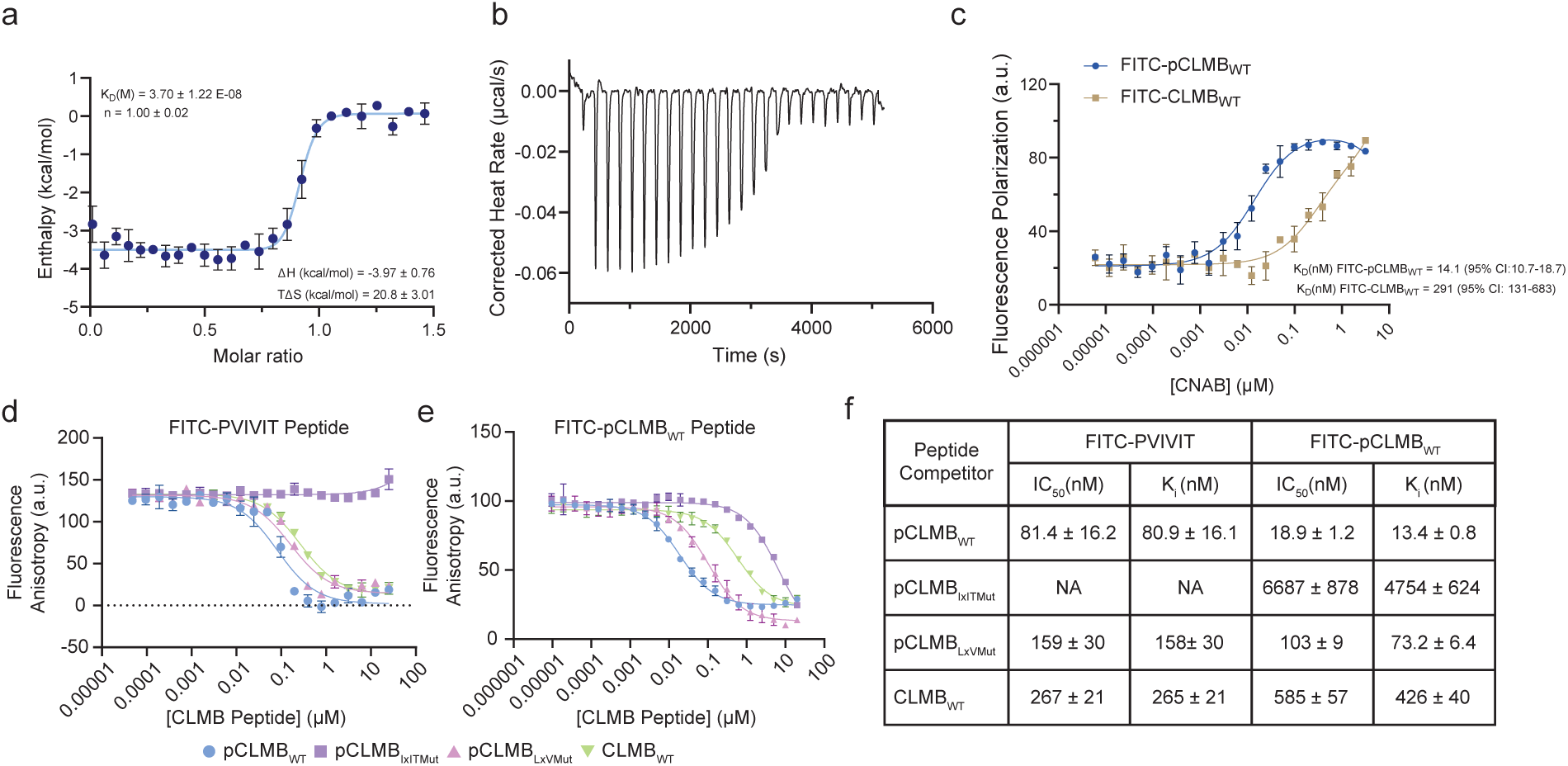
Thr44 Phosphorylation and the LxVP motif contribute to PxIxIT binding. **a.** Determination of the affinity between CNA/B and unlabeled pCLMB_WT_ peptide using isothermal calorimetry. Dissociation constant (*K_D_*) = 37 ± 1.22 nM, reaction stoichiometry (n value) = 1.00 ± 0.02, enthalpy (ΔH) = −3.97 ± 0.76, and entropy (TΔS) = 20.8 ± 3.01 are shown. Data are mean ± s.d. for *n* = 3 independent experiments. **b**. Binding isotherm obtained by titration of unlabeled pCLMB_WT_ peptide into CNA/B (40 μM). **c**. Fluorescence polarization (FP) saturation binding curves of FITC-pCLMB_WT_ and FITC-CLMB_WT_ (5nM) incubated with serial dilutions of CNA/B (0 to 5μM). FP data were fitted to a one-site binding model (mean ± s.d., n = 2 independent experiments). **d**. Competitive binding assay of unlabeled pCLMB_WT_, CLMB_WT_, pCLMB_IxITMut_, and pCLMB_LxVMut_ peptides titrated against FITC-PVIVIT peptide (5 nM) in the presence of 2 μM CNA/B. Data are mean ± s.d. of n = 2 independent experiment. **e.** Competitive binding assay of unlabeled pCLMB_WT_, CLMB_WT_, pCLMB_IxITMut_, and pCLMB_LxVMut_ peptides titrated against FITC-pCLMB_WT_ peptide, (5 nM) in the presence of 0.08 μM CNA/B. Data are mean ± s.d. for n = 2 independent experiments**. f.** Table displaying the experimentally obtained IC_50_ values and converted K_i_ values from data in (**d, e**).

### NMR analysis shows CLMB binding to two sites on calcineurin

Next, to investigate the structure of CLMB in the presence and absence of calcineurin, we used solution-state NMR. 88% of non-proline backbone resonances were assigned using a standard set of triple-resonance experiments, utilizing full-length dually ^13^C- and ^15^N-labeled CLMB (76 residues) that was expressed in *E.coli* (Fig. 4a, Supplemental Fig. 4a). To assess CLMB secondary structure, we employed TALOS-N, a computational method that predicts backbone torsion angles from NMR chemical shifts^23^. This revealed that CLMB is disordered overall, but contains two regions with a tendency to form a β-strand (Supplemental Fig. 4b): Residues I38-I40 include the PxIxIT, consistent with binding of this motif-type to CNA as a β-strand ^22^. N-terminal residues, Q12-V15, were also predisposed to localized folding suggesting possible involvement in other binding interactions. Similar analyses carried out for CLMB using chemical shift secondary structure population interference (CheSPI)^24^ confirmed the absence of secondary structure (Supplemental Fig. 4c).

**Fig. 4.**
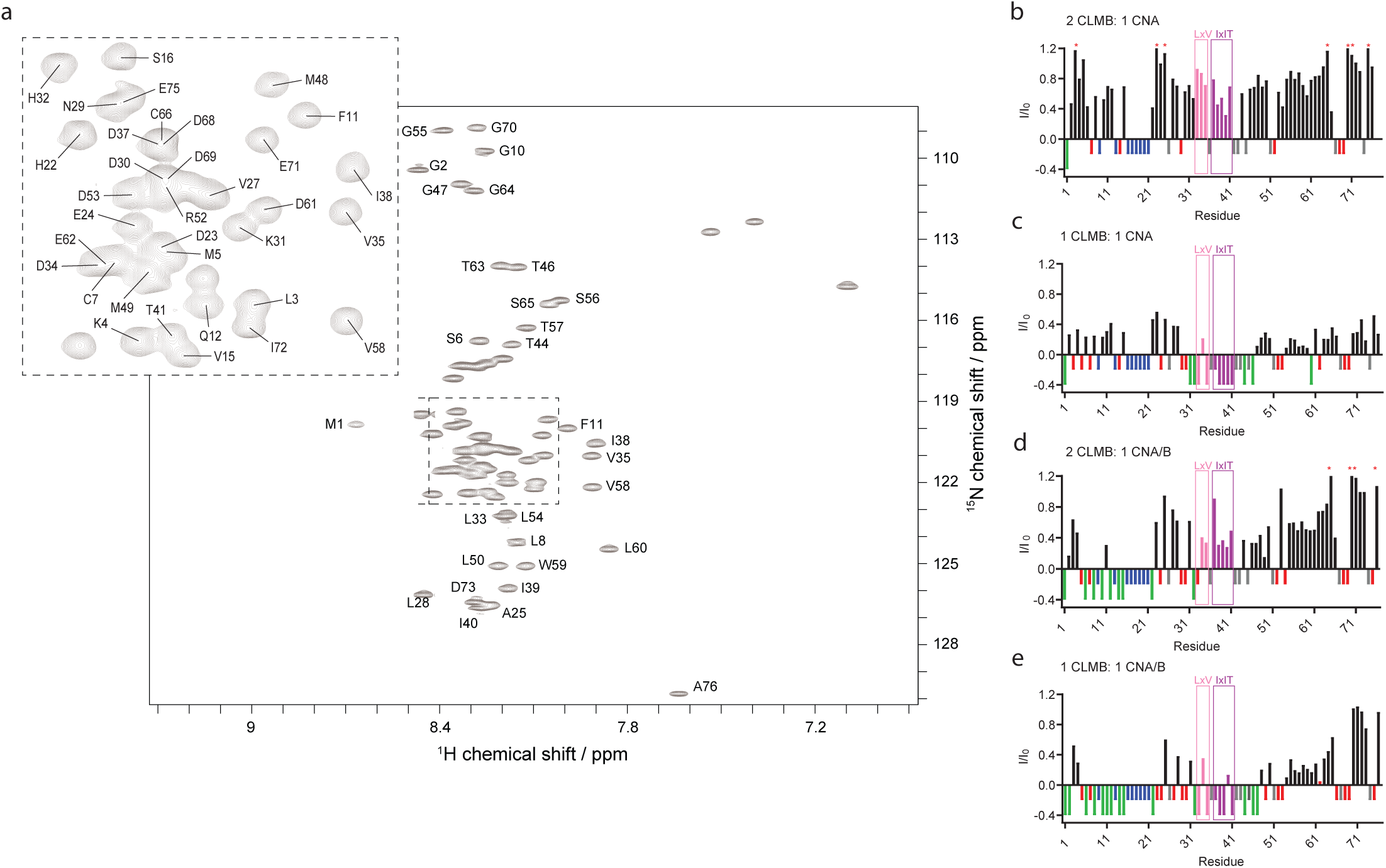
CLMB-calcineurin binding extends beyond the LxVP and PxIxIT motifs. **a.** ^15^N HSQC spectra of free CLMB with the resonance assignment. **b.** NMR spectral analysis of CLMB interaction with CNA at 2:1 (CLMB: CNA); **c**. with CNA at 1:1(CLMB: CNA); **d.** with CNA/B at 2:1 (CLMB: CNA/B); **e**. with CNA/B at 1:1 (CLMB: CNA/B). **b-e.** Graphs display the intensity ratios of CLMB resonances with and without the titrant (CNA or CNA/B) at different molar ratios. The ratios corresponding to the LxV motif (pink) and IxIT (purple) are highlighted. Residues with no access to the intensity ratio are represented with negative numbers, and peaks resulting in complete broadening are shown in green (I/I0 = −0.4). Unassigned residues in apo CLMB are blue (I/I0 = −0.2), prolines (which are not represented in ^15^N-HSQC are grey (I/I0 = −0.2), and residues for which the bound resonance cannot be unambiguously assigned are red, (I/I0 = −0.2). * Residues with I/I0 higher than 1 due to overlapping resonances.

To map the interaction interface of calcineurin on CLMB, we focused first on PxIxIT-mediated interactions by titrating ^15^N-labeled CLMB with varying molar ratios of monomeric recombinant CNA in the absence of CNB and monitoring the interaction through ^15^N-^1^H SOFAST-HMQC experiments (Fig.4). No additional insights were gained from BEST-TROSY experiments^25^. CNA can bind to PxIxIT motifs but lacks the LxVP binding surface found at the interface of CNA and CNB^4^. Due to the size of the CLMB-CNA complex, we anticipated significant broadening of NMR signals at the interaction interface. This broadening can also arise from exchange kinetics between the observable CLMB and the titrant, CNA, making it challenging to distinguish the two effects. As expected, at a molar ratio of 2:1 (CLMB: CNA), many of the CLMB signals undergo significant line broadening (Fig.4b), which become more pronounced at a molar ratio to 1:1, due to the majority of CLMB being bound to CNA (Fig.4c). This is particularly true for signals corresponding to the PxIxIT and LxVP motifs, indicating, as observed in FP experiments, that LxV residues participate in binding to the PxIxIT interface (Fig. 4b, c, Supplemental Fig. 2d). Line broadening across the majority of CLMB residues indicates that the interaction interface on CLMB extends beyond the PxIxIT and LxVP motifs (Fig.4 b, c). Interactions with the N- and C-terminal residues of CLMB may be an avidity-driven effect, anchored by PxIxIT binding, as has been reported for other calcineurin substrates ^26^.

To investigate binding that may occur at the LxVP docking site of calcineurin, we next titrated CLMB with the heterodimeric CNA/B complex (Fig.4d, e). Signal broadening at the N-terminus of CLMB was more pronounced with CNA/B relative to CNA alone, suggesting possible interactions between CLMB and CNB (Fig. 4b, c vs 4d, e, and Supplemental Fig. 4d). These include the Q9-V12 region that displayed partial folding in the absence of calcineurin. At a molar ratio of 2:1 (CLMB: CNA/B), more broadening is observed in LxVP but not PxIxIT residues when compared to the interaction with CNA (Fig. 4d vs 4b, and Supplemental Fig. 4c). This includes not only L33 and V35, but also H32, another residue expected to participate in LxVP binding to CNA/B^7^. However, at a molar ratio of 1:1, where essentially all CLMB should be PxIxIT-bound (given the higher *K*_D_ for PxIxIT vs LxVP and our working concentrations), the pattern of line broadening at the composite motif is similar when complexed with either CNA or CNA/B (Fig. 4c vs 4e). To better visualize LxVP-mediated binding of CLMB to CNA/B, NMR analyses were also performed in the presence of pCLMB_LxVPmut_ peptide (which was not isotopically labeled) to saturate the PxIxIT binding site on CNA/B (Supplemental Fig. 5). While addition of this peptide on its own caused no change in the NMR spectrum for ^15^N-labeled CLMB (Supplemental Fig. 5a), in the presence of CNA/B (2:1 or 1:1 CLMB: CNA) residues in the LxVP motif, particularly H32 and L33, displayed shifting and broadening effects in addition to N-terminal residues, as previously observed (Supplemental Fig. 5b,c). Together these findings show that when in excess (2 CLMB: 1 CNA/B), CLMB engages calcineurin (CNA/B) through both its LxVP and PxIxIT motifs.

To investigate the CLMB-calcineurin interaction further, we carried out structural predictions using AlphaFold 3 for 2 CLMB molecules and CNA/B containing truncated CNA lacking the C-terminal calmodulin binding and regulatory domains (Supplemental Fig. 6a-c) ^27^. Consistent with our NMR analyses, the models show two copies of CLMB simultaneously bound to CNA/B, one each at the LxVP and PxIxIT docking sites. Predicted interactions of pCLMB containing pThr44 with CNA/B was nearly identical, however with the LxVP-bound CLMB molecule showing an additional interaction of pThr44 with the active site, as would occur during catalysis (Supplemental Fig. 6a-c). Thus, these models suggest that formation of multimeric complexes (2 CLMB: 1 CNA/B) will facilitate dephosphorylation of pCLMB.

### Calcineurin dephosphorylates CLMB by forming multimers

Next, we sought to test the prediction that pThr44 dephosphorylation is promoted when CLMB is in excess (2 CLMB: 1 CNA/B) but inhibited at lower CLMB: CNA/B ratios where PxIxIT-bound complexes are favored and protect pThr44 from dephosphorylation. To this end, we inducibly expressed CLMB_WT_-FLAG in HEK293 Flp-In T-REx cells and compared its phosphorylation status with endogenous vs overexpressed calcineurin (i.e. transfection of GFP-CNA, which complexes with endogenous CNB). In stark contrast to a typical substrate, CLMB-FLAG phosphorylation at T44 increased upon calcineurin overexpression (Fig. 5a,b, Supplemental Fig.7a). Intact PxIxIT binding rather than catalytic activity was required for this effect, as overexpressing catalytically inactive calcineurin (containing GFP-CNA_H151Q_) ^28^, but not calcineurin that is defective for PxIxIT binding (containing GFP-CNA_NIR_) ^29^, increased CLMB-FLAG phosphorylation (Supplemental Fig. 7b). We predicted that further increasing CLMB levels in these calcineurin overexpressing cells would promote dephosphorylation of CLMB-FLAG by increasing the CLMB:CNA/B ratio. Thus, we overexpressed V5-tagged CLMB in HEK293 Flp-In T-REx cells that inducibly expressed CLMB-FLAG and overexpressed calcineurin (GFP-CNA complexed with endogenous CNB) (Fig. 5a-c). As predicted, phosphorylation of CLMB_WT_-FLAG decreased upon expression of CLMB_WT_-V5. This effect was dependent on PxIxIT binding, as introducing CLMB_IxITMut_-V5 did not decrease CLMB_WT_-FLAG phosphorylation, but CLMB_LxVMut_-V5, which contains an intact PxIxIT motif, did (Fig, 5a-c). for CLMB_WT_, when CLMB_LxVMut_ was expressed, its phosphorylation was enhanced by overexpression of calcineurin and rescued by co-expression of CLMB_WT_-V5 or CLMB_LxVMut_-V5 but not CLMB_IxITMut_-V5. However, because dephosphorylation is impaired by mutation of the LxV (Fig. 2e), CLMB_LxV_-FLAG displayed increased levels of phosphorylation relative to CLMB_WT_ under all conditions (Fig.5a, b). Finally, we repeated these analyses using CLMB_IxITMut_-FLAG, which displayed partial phosphorylation regardless of calcineurin levels. Surprisingly, CLMB_IxITMut_-FLAG was predominantly dephosphorylated when co-expressed with CLMB_WT_-V5 or CLMB_LxVMut_-V5 but not CLMB_IxITMut_-V5 (Fig. 5a, b). Thus, although the LxVP and PxIxIT components of the CLMB composite motif are both required for efficient dephosphorylation, they do not need to be on the same molecule. A single copy of CLMB must have an LxVP to mediate its own dephosphorylation but can utilize the PxIxIT of neighboring copies to recruit calcineurin, promote LxVP binding, and be dephosphorylated (Fig. 5c, bottom row).

**Fig. 5.**
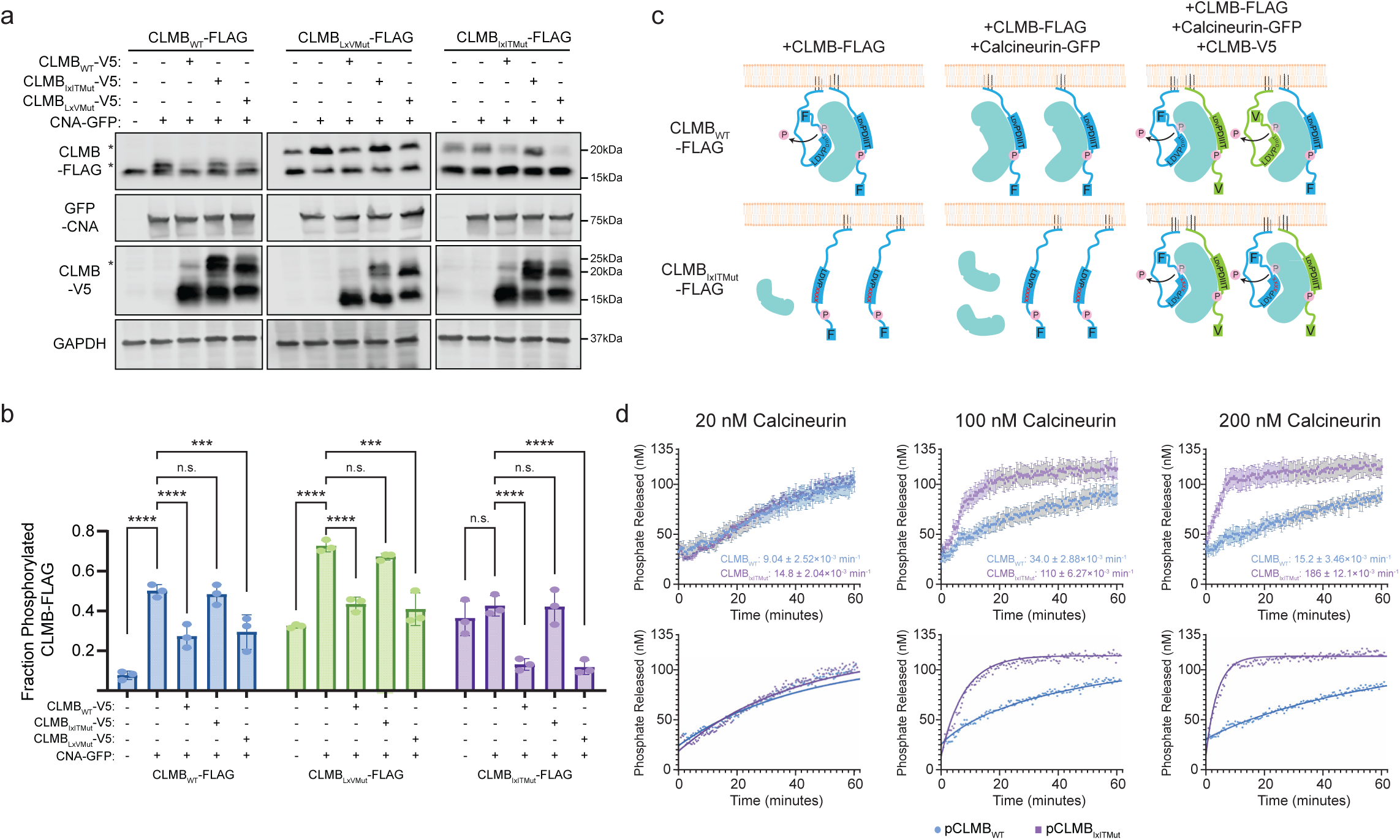
Calcineurin dephosphorylates calcimembrin by forming multimers. **a.** Representative immunoblot showing phosphorylation-induced mobility shifts in CLMB-FLAG (wild-type or indicated mutants) when co-expressed with GFP-CNA and CLMB-V5 (wild-type or indicated mutants). GAPDH was used as a loading control **b.** Data from (**a**) showing mean ± SEM phospho-CLMB-FLAG normalized to total CLMB-FLAG. p-values calculated by two-way ANOVA corrected by Dunnett’s multiple comparison test (n = 3 independent experiments). n.s., not significant; without vs with CLMB_IxIxITmut_-V5 expression: p = 0.995, 0.748, 0.999 from left to right. CLMB_IxITmut_ without vs with CNA-GFP p = 0.626. **** p < 0.0001, *** p < 0.001, p = 0.0005. **c.** Cartoon representation of calcineurin (turquoise)-CLMB complex formation and CLMB dephosphorylation (removal of pink “P”) under conditions in (**a**), i.e. CLMB-FLAG (blue with “F”) overexpression with or without co-expression of Calcineurin-GFP and CLMB-V5 (green with “V”) as indicated. Top row shows CLMB_WT_-FLAG, bottom row shows CLMB_IXITMut_-FLAG (labelled LDVPxxxx). **d**. (Top) Experimental data showing phosphate released from 100 nM pCLMB_WT_ (blue) or pCLMB_IxITMut_ (purple) phosphopeptide dephosphorylated by 20 nM (left), 100 nM (middle) or 200 nM (right) CNA/B and 1 μM calmodulin. Each dephosphorylation rate is displayed and data is mean ± SEM for ≥ 5 replicates at each timepoint. (Bottom) Overlay of experimental data (circles) with results of kinetic simulations (solid lines) using the model shown in Fig. S5a, b for each calcineurin concentration.

To demonstrate more precisely how the ratio of CLMB to calcineurin affects dephosphorylation, we conducted a series of *in vitro* dephosphorylation assays using a fixed substrate concentration (100nM pCLMB_WT_ or pCLMB_IxITMut_ peptides) with increasing concentrations of purified calcineurin (CNA/B) (20, 100 or 200nM) (Fig. 5d, top row). Under these *in vitro* conditions, pCLMB_IxITMut_ was efficiently dephosphorylated due to high enzyme and substrate concentrations and the absence of any competing PxIxIT-containing proteins. For pCLMB_IxITMut_, we observed that dephosphorylation rates increased significantly with the calcineurin concentration (Fig. 5d, top row). In contrast, dephosphorylation of pCLMB_WT_ increased modestly as calcineurin increased from 20 vs 100 nM, and then decreased slightly at 200nM calcineurin, where enzyme concentration exceeded that of substrate (100 nM) (Fig. 5d, top row). To better understand this unusual reaction behavior, we turned to kinetic modeling. First, we developed a model describing the complex series of reactions we expect to occur, based on our NMR and in cell analyses of CLMB-calcineurin interactions (Supplemental Fig. 8). For pCLMB_IxITMut_, which contains only an LxVP motif (Supplemental Fig. 8b), substrate binds to the LxVP-docking site of calcineurin (k_2_, k_-2_) and is dephosphorylated (k_3_) leading to product formation. Thus, dephosphorylation accelerates with increasing enzyme concentration (Fig. 5d, top row). However, for dephosphorylation of pCLMB_WT_ (Supplemental Fig 8a), we propose competition between two binding modes: high affinity binding to the PxIxIT-docking site (k_1_, k_-1_), which prevents pT44 phosphorylation, or lower affinity LxVP-mediated binding (k_2_, k_-2_), which leads to dephosphorylation (k_3_). Furthermore, the model allows one enzyme molecule to bind substrate at each of these sites simultaneously (ES1S2). For this complex reaction mechanism, increasing enzyme concentration drives formation of more LxVP-bound substrate and dephosphorylation, but this is offset by reduced substrate availability due to increased PxIxIT-mediated substrate binding and pThr44 protection. Thus, dephosphorylation rates decrease as enzyme concentration exceeds that of substrate (Fig. 5d, top row).

To further probe this proposed mechanism, we then used kinetic modeling software (KintekExploer^TM^) which fits experimental data directly through dynamic simulation and does not require simplifying assumptions^30^. After providing initial values for rate constants consistent with published reports^6, 31^, our experimental data were fit to the proposed kinetic model (Fig. 5d bottom row and Supplemental Fig. 8). The resulting rate parameters were well within expected values, and the predicted K_D_ for phosphorylated PxIxIT binding to calcineurin was 7nM (Supplemental Fig. 8c), which agrees well with experimentally determined values (Fig. 3a, c). However, we noted that some parameters, particularly on and off rates for substrate binding were poorly constrained by the model. Therefore, we manually manipulated different parameters to better understand how relationships between them influenced the fit to our data (Supplemental Fig. 9). This revealed that the equilibrium binding constant for phospho-PxIxIT binding was critical rather than specific on and off rates (*k_1_* and *k_-1_*) (Supplemental Fig. 9a and b). Furthermore, the *k_1_*and *k_-1_* values predicted by kinetic simulations represented lower limits for these rates as decreasing them caused significant deviations from the data despite keeping the K_D_ constant (Supplemental Fig. 9b). Similarly, *k_off_* for LxVP binding (*k_-2_*) and *k_cat_* (*k_3_*) could be increased proportionally without significantly affecting the overall fit (Supplemental Fig. 9c), suggesting that the catalytic efficiency of LxVP-mediated dephosphorylation is a key driver of the observed reaction kinetics. Confirming this, we found that maintaining the second order rate constant was critical for good agreement between the data and the model (Supplemental Fig. 9d). Thus overall, kinetic modeling of our *in vitro* dephosphorylation data supports the hypothesis that phosphorylated CLMB is dephosphorylated via multimer formation. This novel mechanism, which requires substrate levels to exceed that of calcineurin for efficient dephosphorylation, is dictated by the composite LxVPxIxIT motif.

## Discussion

Here we describe multivalent interactions between calcineurin and CLMB, an intrinsically disordered microprotein and calcineurin substrate that dynamically associates with membranes via protein lipidation. We elucidate roles for a novel composite motif in CLMB, LxVPxIxIT, that combines two SLiMs, PxIxIT and LxVP, with distinct binding modes and functions in calcineurin signaling.

Calcineurin, the conserved and ubiquitously expressed calcium/calmodulin regulated protein phosphatase, has been categorized as a dynamic or date hub and specifically as a linear motif binding hub (LMB-hub) because it engages a diverse set of partners that bind competitively to the structurally plastic PxIxIT docking groove^32^. PxIxIT peptides from natural substrates bind to calcineurin as a β strand, vary greatly in sequence and typically have low affinities (∼2.5 to 250 micomolar)^26, 33^, while regulators and scaffolds, including PVIVIT, the selected inhibitor peptide, have K_D_’s of ∼0.5 micromolar ^21, 34, 35^. By contrast, we show that the ^32^HLDVPDIIITPP(p)TP^45^ sequence in CLMB binds calcineurin at the PxIxIT-docking groove with a K_D_ of ∼14 nM. This extraordinary affinity is mediated predominantly via binding of the core PxIxIT motif which is enhanced ∼20 fold by phosphorylation at the +3 position (pThr44), and ∼5 fold by the N-terminal LxV sequence (Fig. 3). Such high affinities have also been achieved by engineering the PVIVIT peptide to include a hydrophobic residue at the −1 position and a phosphothreonine at the +3 position, although phosphorylation does not universally enhance PxIxIT affinity^35^. A natural PxIxIT peptide derived from an endogenous inhibitor of calcineurin (CABIN1),^2145^PPEITVTPP(p)TP^2156^ also shows high affinity (∼170 nM) due to phosphorylation of its C-terminal sequence, PPTP, which is identical to that found in CLMB^34^. This proline-rich sequence is predicted to form a polyproline helix that may function to extend the β-strand interaction with calcineurin^36, 37^. Thus, the composite LxVPxIxIT motif in CLMB combines several features that consistently stabilize PxIxT-docking. Because calcineurin is an LMB-hub, such high affinity PxIxIT peptides typically inhibit signaling by preventing its interaction with substrates^35, 38^. It is surprising, therefore that CLMB exhibits high affinity PxIxIT-mediated docking yet is also dephosphorylated by calcineurin. As we demonstrate, this is only possible due to the LxVP portion of the composite SLiM in CLMB, and dephosphorylation can only occur when excess CLMB levels allow lower affinity LxVP-docking to occur on a molecule of calcineurin whose PxIxIT-docking site is saturated.

Based on our findings, we present two models for how CLMB may shape calcineurin activity *in vivo* (Figure 6). First, dynamic S-acylation promotes CLMB association with membranes (Fig. 6a, Step 1), where, when phosphorylated, it recruits a localized pool of calcineurin via high affinity PxIxIT-mediated binding, (Fig. 6a, Step 2). This membrane sequestration may prevent calcineurin from dephosphorylating substrates in other cellular locations. In the presence of a Ca^2+^ signal, Ca^2+^ and CaM then activate calcineurin, which exposes the LxVP docking site^39^. If CLMB is in excess relative to calcineurin and PxIxIT binding is fully saturated, a second CLMB molecule will bind via its LxVP motif which positions pThr44 for dephosphorylation at the active site (Fig. 6a, Step 3). However, as the population of CLMB shifts to the dephosphorylated form, PxIxIT binding affinity decreases (Fig. 3) which promotes calcineurin dissociation from CLMB, allowing the enzyme to engage and dephosphorylate other nearby LxVP- and PxIxIT-containing substrates (Fig. 6a, Step 4). This tuning of PxIxIT affinity is critical for signaling as demonstrated for AKAP5, a scaffold that concentrates both calcineurin and NFAT near L-type Ca^2+^ channels in neurons to promote NFAT dephosphorylation^40^. In that case, increasing the AKAP5 PxIxIT affinity for calcineurin ∼two-fold significantly impaired NFAT activation due to increased sequestering of calcineurin^41^.

**Fig. 6.**
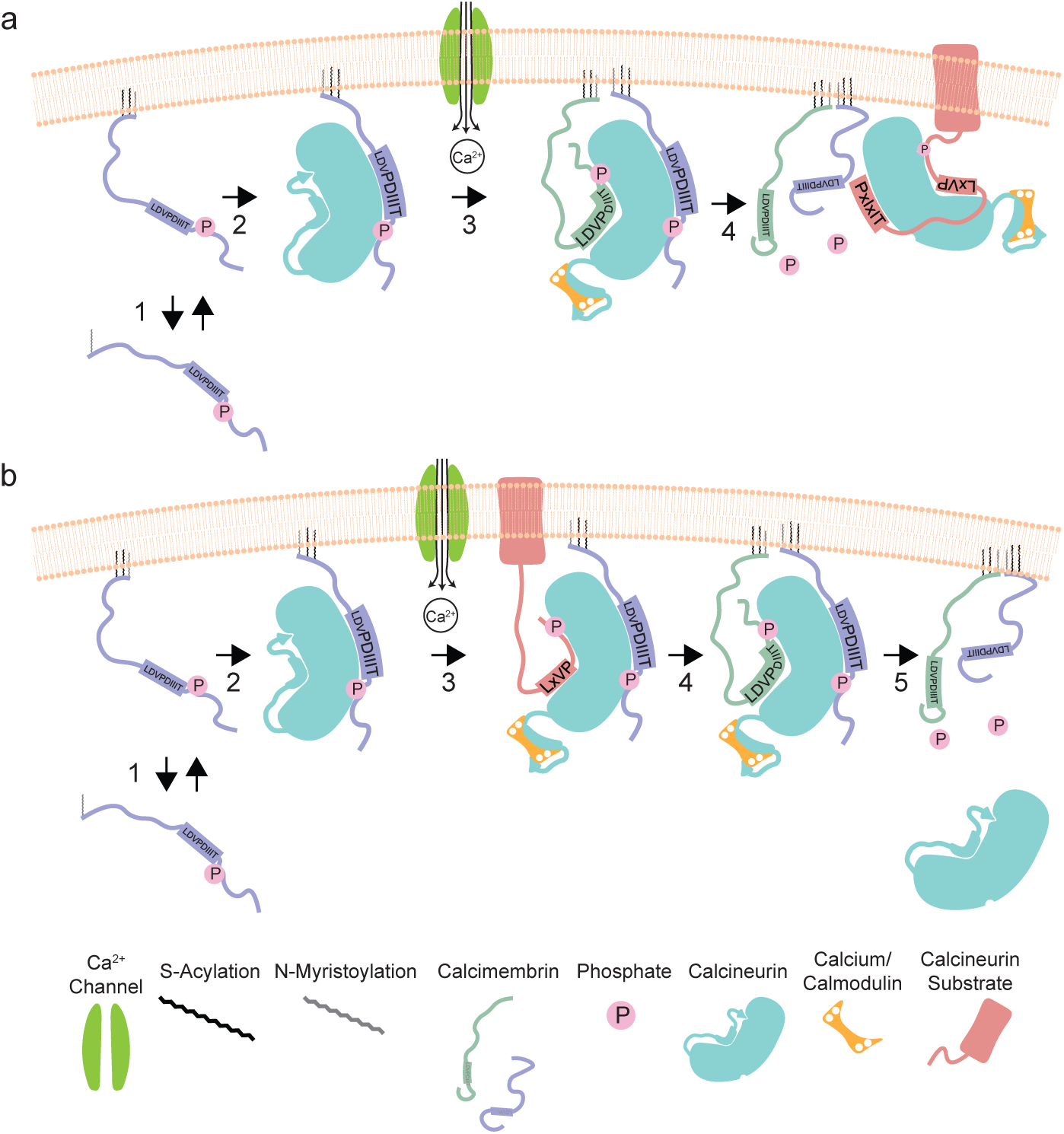
Models explaining how calcimembrin may modulate calcineurin signaling. **a** (1) CLMB localizes to membranes via dynamic acylation where it (2) recruits calcineurin (turquoise). At low CLMB levels, calcineurin remains tightly bound to phosphorylated CLMB (purple) regardless of Ca^2+^ signals. (3) At higher CLMB levels, CLMB (purple and green) forms multimers with calcineurin and is dephosphorylated upon calcineurin activation by Ca^2+^ and CaM. (4) Calcineurin is then released and can engage and dephosphorylate membrane-associated substrates (red) that contain PxIxT and LxVP SLiMs. **b** (1) CLMB localizes to membranes via dynamic acylation where it (2) recruits calcineurin (turquoise). At low CLMB levels, calcineurin remains tightly bound to phosphorylated CLMB (purple). (3) Ca^2+^ and CaM activate calcineurin, allowing it to engage and dephosphorylate LxVP-containing substrates (red). (4) At high CLMB levels (purple and green) calcineurin-CLMB multimers form. (5) This promotes CLMB dephosphorylation and triggers calcineurin release to terminate signaling.

In our second model (Fig. 6b), active calcineurin-pCLMB dimers (1 CLMB: 1 CNA/B) directly engage and dephosphorylate substrates at the membrane that contain an LxVP, but no PxIxIT motif (Fig. 6b, step 3). Such substrates are exemplified by PRKAR2A, the type II-alpha regulatory subunit of protein kinase A, which contains an LxVP but depends on the PxIxIT motif within its binding partner, AKAP5, to recruit calcineurin and promote dephosphorylation^9, 42^. Proteome-wide predictions suggest that many proteins may similarly harbor an LxVP without an associated PxIxIT ^9^. In our second model (Fig. 6b), active calcineurin-pCLMB dimers (1 CLMB: 1 CNA/B) directly engage and dephosphorylate such PxIxIT-less substrates at the membrane (Fig. 6b, step 3). Simply recruiting calcineurin to membranes, where association kinetics are 20-30 fold faster than in the cytosol of eukaryotic cells, may enhance LxVP binding causing their efficient dephosphorylation in the absence of a PxIxIT motif ^43^. Such division of the PxIxIT and LxVP motifs among two molecules may also allow for the extremely high affinity PxIxIT observed in CLMB. In substrates that contain both a LxVP and PxIxIT motif, stable PxIxIT binding would be expected to limit enzymatic efficiency by slowing product release. However, acting only as a scaffold, CLMB can remain tightly associated with calcineurin via PxIxIT-binding while other substrates cycle in and out of the LxVP-binding surface to be dephosphorylated. In this scenario (Fig. 6b), calcineurin remains at the membrane either until CLMB levels increase sufficiently to allow CLMB dephosphorylation and calcineurin dissociation (Fig. 6b step 4) or until CLMB is de-acylated. In both models, intracellular Ca^2+^ dynamics, CLMB protein levels, and S-acylation/de-acylation activities combine to modulate calcineurin signaling. Future studies aim to identify the palmitoyl acyltransferases (PATs) and acyl protein thioesterases (APTs) that dynamically regulate CLMB S-acylation to better understand how these critical regulators contribute to calcineurin-CLMB signaling *in vivo*.

Another missing player in our model is the kinase(s) that phosphorylates CLMB at Thr44. One candidate is the stress-activated p38 mitogen-activated protein kinase (MAPK), which phosphorylates the identical PxIxIT-flanking sequence in Cabin1 (PP(p)TP)^44^. Because phosphorylation increases CLMB-calcineurin affinity, kinase activation is required to prime regulation of calcineurin signaling by CLMB. However, phosphorylation also establishes a negative feedback mechanism, with dephosphorylation decreasing calcineurin’s interaction with its substrate. Thus, regulators that modify kinase activity may also significantly affect CLMB-calcineurin signaling dynamics.

This study highlights the critical role that SLiMs play in conferring flexibility, tunability and dynamism to cell signaling at the molecular level. Further studies are needed to establish whether composite LxVPxIxT motifs always limit substrate dephosphorylation to occurring in a multimeric complex. Of at least 30 human proteins containing predicted LxVPxIxIT motifs, several are ion channels, such as TRPM6, that assemble into higher order complexes^45^, suggesting that these motifs could be a fidelity mechanism that limits dephosphorylation to substrates that are properly assembled into complexes. Similarly, multivalency is a characteristic exhibited by many SLiMs. Some composite SLiMs bind multiple partners in an exclusive manner, for example requiring a kinase and a phosphatase to compete for access to a shared substrate^46, 47^. For the intermediate chain of dynein, multivalency regulates assembly of the dynein complex and its interaction with distinct binding partners^48^. In the cell cycle regulator CDKN1A (p21), sequences that flank the PCNA-binding SLiM also encode an essential nuclear localization sequence^49^. Thus, ‘composite’ SLiMs that encode multiple interactions may be more common that currently appreciated. This complexity emphasizes how critical SLiM-based interactions are for integrating and coordinating the plethora of possible protein-protein interactions that dictate cellular responses.

A major challenge that lies ahead is to identify the molecular targets and physiological functions that are regulated by calcineurin-CLMB. While this manuscript was being revised for publication, another study showed that C16orf74/MICT1 is highly expressed in brown fat in mice and controls thermogenesis by regulating the dephosphorylation of PKA RIIβ by calcineurin^50^. In some pancreatic ductal adenocarcinoma (PDAC) derived cell lines, CLMB increases proliferation and migration suggesting that CLMB shapes calcineurin signaling to promote oncogenesis^10^. Indeed, high CLMB expression is associated with poor patient outcomes for PDAC, melanoma and head and neck cancer^10, 51^. Mechanisms underlying these activities have not yet been identified, but these cancers may provide good starting points for examining how CLMB modifies calcineurin signaling in a clinically relevant context.

## Methods

### Cell culture and transfection

Cells were cultured at 37 °C in 5% CO_2_. HEK293 Flp-In TReX and MCF7 cell lines were grown in standard media containing Dulbecco’s modified eagle’s medium (DMEM) with 4.5 mg/mL glucose, L-glutamine, and sodium pyruvate (Cytiva SH30243.FS) supplemented with 10% fetal bovine serum (Corning MT35010CV). Transfections were done according to manufacturer instructions. HEK293 cells were transfected using lipofectamine 2000 (Thermo Fisher Scientific 11668019). MCF7 cells were transfected using Transfex ATCC transfection reagent (ATCC ACS-4005).

### Stable cell line generation

On day one, 9.6 x 10^5^ HEK293 Flp-In TReX cells were transfected using lipofectamine 2000 with 1.5 µg pOG44 and 0. 5 µg pcDNA5 FRT TO encoding proteins of interest in a 6 well dish (plasmid # 2801, 2852, 2860, 2861, 2862, 2863, 2864, and 2880). Transfections were done according to manufacturer instructions. On day two, transfected cells were passaged onto 10cm plates. On day three, selection was started using media containing 3 µg blasticidin and 200 ng/mL hygromycin. Selection media was replaced every 2-3 days until distinct colonies appeared. Colonies were selected using cloning rings and expanded before being stored in liquid nitrogen.

### Metabolic labeling to detect CLMB lipidation

HEK293 Flp-In TReX lines stably expressing CLMB_WT_, CLMB_C7,14S_, or CLMB_G2A_ were induced with 10 ng/mL doxycycline for 24 hours and split evenly among three 10 cm tissue culture dishes. The following day, cells were rinsed with PBS and incubated in DMEM containing 4.5 mg/mL glucose, L-glutamine, sodium pyruvate, 2% FBS, and either 20 µM YnMyr (Click Chemistry Tools 1164-5), 30 µM 17-ODYA (Cayman Chemical Company 34450-18-5), or vehicle (ethanol and DMSO) overnight. Cells were then co-incubated with 20 µM YnMyr, 30 µM 17-ODYA, or vehicle and 20 µM MG132 (EMD Millipore 474787-10MG) for 4 hours before washed twice in PBS and frozen at −80 °C. Cells were lysed in 400 µL TEA lysis buffer (50 mM TEA pH 7.4, 150 mM NaCl, 1% triton X-100, and 1 mM PMSF) by inverting for 15 minutes at 4 °C. Lysates were passed five times through a 27-gauge needle, centrifuged at 16,000 x *g* for 15 minutes at 4 °C and protein content determined by Pierce BCA protein assay kit (Thermo Fisher Scientific 23225). 25 µg total lysate was removed as input and 0.5 mg total protein was incubated with pre-washed magnetic M2 anti-FLAG (Sigma-Aldrich M8823-5ML) beads at 4 °C for 1.5 hours while inverting. Beads were washed 3X with modified RIPA (50 mM TEA pH 7.4, 150 mM NaCl, 1% Triton X-100, 0.1% SDS, and 1% sodium deoxycholate) and resuspended in 40 µL 1x PBS. 10 µL of 5x click reaction mix (1x PBS, 0.5 mM biotin azide, 5 mM TCEP, 0.5 mM TBTA, 5 mM CuSO_4_) was added and reactions incubated for 1 hour at 24 °C, shaking at 550 RPM. Beads were washed 3X in modified RIPA and eluted 2X using 30 µL of 1 mg/mL FLAG peptide in modified RIPA buffer. 12 µL of non-reducing 6x SDS sample buffer was added to eluate before heating at 95 °C for 2 minutes and analysis by SDS-PAGE. Note that addition of reducing agents to SDS sample buffer results in loss of 17-ODYA signal. Samples were transferred to nitrocellulose membrane and lipidation was detected using fluorophore-tagged streptavidin, and total CLMB with anti-FLAG. Immunoblots were imaged using a Licor Odyssey CLx and quantified with Image Studio software.

### Metabolic pulse-chase labeling assay to assess dynamic CLMB S-acylation

HEK293 Flp-In TReX cells stably expressing CLMB_WT_-FLAG were induced with 10 ng/mL doxycycline for 48 hours. Cells were rinsed 2X in PBS and incubated in methionine free DMEM supplemented with 5% charcoal-stripped FBS, 1 mM L-glutamate, 1 mM sodium pyruvate, 0.4 mM cysteine and 10 ng/mL doxycycline along with vehicle (ethanol), 30 µM 17-ODYA, 50 µM L-AHA, or 17-ODYA and L-AHA together. After two hours, cells were washed 2X in PBS and either harvested immediately or incubated in standard media supplemented with 10 µM palmitic acid and 10 ng/mL doxycycline, with or without 10 µM palmostatin B (EMD Millipore 178501-5MG). Cells were rinsed in PBS and frozen at −80 °C. Metabolic labeling was done as described in the previous section with the exception that two rounds of labeling were done. First, Cy5-azide (Click Chemistry Tools AZ118-1) was linked to 17-ODYA. Then, cells were washed and L-AHA was linked to AF-488 (Click Chemistry Tools 1277-1) alkyne before being washed three additional times and eluted using FLAG peptide. Samples were resolved via SDS-PAGE and in-gel fluorescence was read using the GE Amersham Typhoon imaging system. Images were quantified in Image Studio. The proportion of S-acylated CLMB is calculated as Cy5/A-488 normalized to the no chase control.

### Acyl-PEG exchange

HEK293 Flp-In TReX lines stably expressing CLMB_WT_, CLMB_C7S_, CLMB_C7,14S_, or CLMB_G2A_ were induced with 10 ng/mL doxycycline for 48 hours. Cells were rinsed 2X in PBS and stored at −80 °C. Cells were lysed in 400 µL lysis buffer (50 mM TEA pH 7.3, 150 mM NaCl, 2.5% SDS, 1 mM PMSF, 100 units benzonase) at 37 °C for 20 minutes, shaking at 800 RPM. Lysates were passed 10X through a 27-gauge needle, centrifuged at 16,000 x *g* for 5 minutes and protein quantified by BCA. Lysates were adjusted to 2 mg/mL and incubated with 10 mM TCEP for 30 minutes at room temperature. Cysteines were protected by incubating with 25 mM NEM for 2 hours at 40 °C, shaking at 800RPM. Protein was precipitated over night with four volumes of acetone at −20 °C, centrifuged at 16,000 x *g* for 20 minutes at 4 °C and washed 4X in 70% acetone. Acetone was removed, pellets air-dried to remove all traces of acetone and resuspended in 75 µL Buffer A (50 mM TEA pH 7.3, 150 mM NaCl, 4% SDS, 0.2% triton X-100, 4 mM EDTA) for 1 hour at 40 °C, shaking at 800 RPM. Samples were centrifuged at 16,000 x *g* for 2 minutes, 7 µL removed as input, and the remainder split into +cleavage and −cleavage fractions. The +cleavage sample containing 1% SDS was treated with 750 µM hydroxylamine (final concentration) for 1 hour at 30 °C, rotating end over end. Following methanol-chloroform precipitation (ice cold, final concentrations 42% methanol, 16% chloroform) samples were centrifuged at 16,000 x *g* for 20 minutes at 4 °C, aqueous layer removed, washed 3X with 100% methanol and air dried. Pellets were resuspended as described in Buffer A and incubated in 1 mM mPEG-Mal (Methoxypolyethylene glycol maleimide, 5 kDa, 63187 Sigma-Aldrich) for 2 hours at room temp, rotating end over end. Samples were methanol-chloroform precipitated as described, resuspended in Lysis buffer with 1X SDS-PAGE sample buffer, resolved using SDS-PAGE and visualized via immunoblotting with mouse anti-FLAG to detect CLMB-FLAG. Endogenous calnexin was detected using anti-calnexin antibodies. Immunoblots were imaged using a Licor Odyssey CLx and quantified with Image Studio software.

### Detergent-assisted subcellular fractionation

HEK293 Flp-In TReX lines stably expressing CLMB_WT_, CLMB_C7,14S_, or CLMB_G2A_ were induced with 10 ng/mL doxycycline for 48 hours and 20 µM MG132 (EDM Millipore 474787-10MG) for 4 hours prior to harvesting. Cells were washed 2X in PBS, harvested and immediately lysed in 600 µL of Cytoplasm Buffer (10 mM HEPES pH 6.8, 100 mM NaCl, 300 mM sucrose, 3 mM MgCl_2_, 5 mM EDTA, 0.015% digitonin, 1x halt protease inhibitor cocktail (Thermo Fischer scientific 78442)) at 4 °C for 15 minutes. 60 µL of total lysate was removed as input and the remainder centrifuged at 1000 x *g* for 10 min at 4 °C. The supernatant was removed as cytoplasmic fraction and the pellet washed 2X in PBS and resuspended in 600 µL membrane buffer (10 mM HEPES pH 7.5, 100 mM NaCl, 300 mM sucrose, 3 mM MgCl_2_, 3 mM EDTA, 0.5% Triton X-100, 1x halt protease inhibitor cocktail) for 30 minutes at 4 °C turning end over end. Insoluble material was pelleted by centrifugation at 5000 x *g* for 10 minutes at 4 °C and supernatant removed as membrane fraction. Both fractions were centrifuged at 12,000 x *g* at 4 °C for 10 minutes. Samples were heated in 1X SDS sample buffer at 95 °C for 5 minutes, resolved via SDS-PAGE, immunoblotted and CLMB-FLAG detected using anti-FLAG antibodies; anti-GM130 and anti-α-tubulin were used as membrane and cytosolic markers respectively. Immunoblots were imaged using a Licor Odyssey CLx and quantified with Image Studio software. CLMB in cytosolic and membrane fractions were quantified as CLMB_Cyto_/(CLMB_Cyto_+ CLMB_Mem_) and CLMB_Mem_/(CLMB_Cyto_ + CLMB_Mem_), respectively.

### Immunofluorescence, microscopy, and image analysis

MCF7 cells were co-transfected (as described previously) with equal amounts of pcDNA5 encoding CLMB_WT_, CLMB_IxITMut_, CLMB_LxVMut_, CLMB_T44A_ and pcDNA5 encoding CNA-GFP. The following day cells were plated on poly-lysine coated 12 mm, #1.5H glass coverslips (ThorLabs). On day three, cells were washed in PBS and fixed in 100% methanol for 10 minutes at −20 °C. Coverslips were washed 3 times in 1x PBS, blocked (1x PBS, 200 mM glycine, 2.5% FBS) for 30 minutes at room temperature, then incubated in primary antibody diluted in blocking buffer for 1 hour before being washed 3 times in 1x PBS and incubated for 1 hour in secondary antibody solution. Following three additional washes, coverslips were mounted using Prolong Glass mountant with NucBlue stain. After sitting overnight at room temperature, coverslips were imaged using a Lionheart FX automated widefield microscope with a 20X Plan Fluorite WD 6.6 NP 0.45 objective. For each coverslip, a 7 by 7 grid of non-overlapping images was automatically taken. All images were manually assessed and those with defects (debris, bubbles, out-of-focus, etc.) were filtered out. Images were analyzed using an automated pipeline using CellProfiler ^52^. Briefly, cells were identified by their nuclei and all those lacking CLMB-FLAG staining removed. CLMB-FLAG signal was used to define the cell body and all cells containing one or more fully saturated pixels in the CLMB-FLAG or CNA-GFP channels were removed. Pearson’s correlation coefficient between CLMB-FLAG and CNA-GFP was determined for each individual cell. >500 cells per condition were analyzed.

### GFP-Trap coimmunoprecipitation

HEK293 cells were co-transfected (as described previously) with pcDNA5 encoding CNA-GFP and pcDNA5 encoding CLMB_WT_, CLMB_IxITMut_, CLMB_LxVMut_, CLMB_T44A_, or CLMB_LxVMut+T44A_. 24 hours post-transfection, cells were washed 2X in PBS and frozen at −80 °C. Cells were lysed in 500 µL lysis buffer (50 mM Tris pH 7.5, 150 mM NaCl, 1% NP-40), aspirated 5X with a 27-gauge needle, centrifuged at 16,000 x *g* for 20 min at 4 °C and protein quantified via BCA. 100 mg total protein was removed as input. Lysates were adjusted to 0.5% NP-40 and 500 µg of protein incubated with pre-washed magnetic GFP-TRAP beads (ChromoTek gtma-20) for 2 hours at 4 °C, turning end over end. Beads were magnetized, washed 3 times and eluted with 50 µL of 2x SDS PAGE sample buffer. Samples were resolved via SDS-PAGE and visualized via immunoblotting with anti-flag antibodies to detect CLMB-FLAG and anti-GFP to detect CNA-GFP. Anti-GAPDH was used as a loading control for inputs. Immunoblots were imaged using a Licor Odyssey CLx and CLMB-calcineurin binding was quantified with Image Studio software. Co-purified CLMB was normalized using the following: (CLMB_IP_/CLMB_Input_)/GFP-CNA_IP_. Additionally, all values are displayed relative to CLMB_WT_ which is set at 1.

### Protein expression and purification

CNA catalytic dead mutant H151Q with an N-terminal His_6_-CNA (residues 1–391) and CNB (residues 1– 170) subunits were expressed using *E. coli* BL21(DE3) cells in Lysogeny Broth (LB) supplemented with 50 μg/mL ampicillin. Protein expression was induced with 1 mM isopropyl β-d-1-thiogalactopyranoside (IPTG) at 18 °C. Cells were lysed using a Fisherbrand™ Model 505 Sonic Dismembrator and lysates were centrifuged at 46,000 × *g* at 4 °C for 45 minutes. The protein was purified by incubation with nickel-nitriloacetic acid (Ni-NTA) agarose resin (Qiagen) for 3h at 4 °C. The resin was washed with 50 mM Tris pH 8.0, 250 mM NaCl, 20 mM imidazole, and 1 mM TCEP. A buffer containing 50 mM Tris pH 8.0, 250 mM NaCl, 300 mM imidazole, and 1 mM TCEP was used for elution. Calcineurin was further purified by SEC (Superdex® 200 Increase 10/300 GL) using Phosphate-buffered saline (PBS) and CaCl_2_ 1mM as the final buffer.

### Fluorescence polarization binding assay

All FP measurements were performed on a multifunctional microplate reader (EnVision, Perkin Elmer) in black 384-well microplates (Corning, Cat No 3575) with 30 µL of the assay solution per well. 480-nm excitation and 535-nm emission filters were used for the FP measurements. In the FP saturation binding experiments, 5 nM FITC labeled peptides were mixed with increasing concentrations of calcineurin in PBS buffer (pH 7.4) (commercially available tablets, Thermo Fisher #18912014) and 500 µM CaCl_2_ buffer (from 0 to 25 μM). Where indicated (Supplemental Fig. 3), these analyses were carried out at different pHs using a buffer with 20 mM Hepes pH 6.9, 100 mM NaCl, 0.5 mM EDTA, 1 mM TCEP or a buffer containing 50 mM HEPES, pH 8.0, 150 mM NaCl, 1 mM TCEP, 500μM CaCl_2_. FP values were plotted against the log of the protein concentrations, and the dissociation constant (apparent *K*_D_) was obtained from the resulting sigmoidal curve as analyzed in GraphPad Prism 10. Protein concentrations above 5µM resulted in an upturn in fluorescence polarization signal, indicative of possible aggregation or artifacts; to maintain assay integrity, the considered concentrations were limited to ≤5µM.

### Fluorescence polarization competition assay

For the competitive binding assay, a mixture containing 5 nM FITC-labeled CLMB peptides and CNA/B (0.08 and 2 µM for CLMB vs PVIVIT experiments) was incubated with serial dilutions of unlabeled peptides for 30 minutes at room temperature. The FP values were determined, and the IC_50_ values, i.e., the concentrations required for 50% displacement of the FITC-labeled peptide, were calculated in GraphPad Prism 10. The *K*_i_ values of competitive inhibitors were calculated based on a previously reported method^53^, further adapted by^54^. Briefly, the relation of *K*_i_ and IC_50_ in competitive inhibition as applied to protein-ligand–inhibitor (P–L–I) interactions, can be summarized by the following equation:

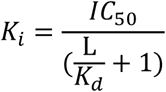

Where L stands for the concentration of ligand, i.e., the FITC-labeled peptide used in the FP binding assay; and *K*_D_ corresponds to the dissociation constant of the same FITC-labeled peptide resulting from the performed FP binding assay.

### ITC measurements

Isothermal titration calorimetry (ITC) experiments were performed using an Affinity ITC from TA Instruments (New Castle, DE) equipped with an autosampler in a buffer containing 50 mM HEPES, pH 8.0, 150 mM NaCl, 1 mM TCEP, 500µM CaCl_2_ at 25 °C. 40 μM protein solution in the calorimetric cell was titrated with 100 μM peptide solution using 0.5 µL injection in 200-sec intervals using stirring speed at 125rpm. The resulting isotherm was fitted with a single site model to yield thermodynamic parameters of ΔH, ΔS, stoichiometry, and *K*_D_ using NanoAnalyze software (TA instruments). All results are representative of three independent experiments (n = 3).

### NMR analyses

All NMR data were acquired using TopSpin3.4 (Bruker), processed with NMRPipe v.2020/2021 and analyzed with CCPNmr v3^55, 56^. Structural biology applications were compiled by the SBGrid consortium. Three-dimensional (3D) NMR experiments were conducted on a Bruker spectrometer operating at 600 MHz, equipped with TCI cryoprobes and z-shielded gradients. Samples of approximately180-μM 2H-, ^13^C- and ^15^N-calcimembrin in 20 mM Hepes pH 6.9, 100 mM NaCl, 0.5 mM EDTA, 1 mM TCEP and 5% v/v ^2^H2O were used. Temperature was set to 25°C. Chemical shift assignments are deposited in the Biological Magnetic Resonance Bank (BMRB) under accession code 52641.

The NMR titration samples were prepared by adding unlabeled calcineurin A/B complex or unlabeled calcineurin subunit A into solutions of U-[^15^N]-labeled calcimembrin (50 μM) to reach a ratio of 0.5:1 (25 μM unlabeled calcineurin) or 1:1 (50 μM unlabeled calcineurin). Titration experiments featuring pCLMB_LxVMut_ (40µM) peptide with free CLMB (50µM), and with calcineurin:CLMB at 1:1 and 0.5:1 stoichiometries were also performed. Two-dimensional (2D) spectra were acquired using SOFAST-HMQC on a Bruker 800 MHz Avance III spectrometer equipped with a TXO cryoprobe and z-shielded gradients^57^. Data was processed using NmrPipe and analyzed with CCPNmr. Secondary structure prediction was analyzed using TALOS-N^58^ and CheSPI^24^.

### AlphaFold 3 modeling

Calcineurin-CLMB complex modeling was done using AlphaFold 3 on AlphaFold server^27^. Modeling was done using truncated CNA_1-391_, CNB, and two copies of CLMB, either unmodified or phosphorylated at Thr44. Zn^2+^ and Fe^3+^ (found in the calcineurin active site) and four Ca^2+^ ions (bound to CNB) were included in modeling. All images were generated in UCSF ChimeraX 1.8^59^.

### *In vivo* analysis of CLMB phosphorylation status

HEK293 FLIP-In TreX cells stably expressing CLMB_WT_-FLAG, CLMB_IxITMut_-FLAG, CLMB_LxVMut_-FLAG, or CLMB_T44A_-FLAG were transfected with empty vector or pcDNA5 FRT TO encoding CNA-GFP alongside empty vector or pcDNA5 FRT TO encoding CLMB_WT_-V5, CLMB_IxITMut_-V5, or CLMB_LxVMut_-V5. Protein expression was induced for 24 hours with 10 ng/mL doxycycline, cells washed 2X in PBS and stored at −80 °C. Cells were lysed (50 mM Tris pH 8, 150 mM NaCl, 1% NP-40, 0.1% SDS, 0.5% sodium dodecyl sulfate, 1x Halt protease inhibitor cocktail) and resolved via SDS-PAGE and detected via immunoblotting.

### *In vitro* CLMB phosphopeptide dephosphorylation

CLMB_WT_, CLMB_IxITMut_, and CLMB_LxVMut_ phospho-peptides were synthesized by Vivitide, resuspended in 10 mM Tris pH 8.0 and dephosphorylated at 26 °C in reaction buffer (50 mM Tris pH 8.0, 100 mM NaCl, 6 mM MgCl_2_, 2 mM CaCl_2_, 1 mM EGTA, 0.1 mg/mL BSA, 1 mM DTT, 0.5 µM MDCC-PBP (Thermo Scientific PV4406), 1 µM CaM (Millipore Sigma 208694-1MG)). Reactions were initiated by addition of CLMB peptide to a final concentration of 100nM. Reactions took place in a total of 50 µL in a corning black 96 well vessel and were monitored continuously via fluorescence emitted by phosphate bound MDCC-PBP (EX:425/ EM:460). Phosphate standards were used to convert fluorescence intensity to phosphate concentration. Data were analyzed using Prism software. Because the experimental design does not comply with assumptions of standard kinetic models, reaction rates were obtained by fitting data to a single exponential model as an empirical description of the curves to facilitate comparison between experimental conditions.

### *In vitro* RII phospho-peptide dephosphorylation

1 µM RII peptide was dephosphorylated at 26 °C in reaction buffer (50 mM Tris pH 8.0, 100 mM NaCl, 6 mM MgCl_2_, 2 mM CaCl_2_, 1 mM EGTA, 0.1 mg/mL BSA, 1 mM DTT, 0.5 µM MDCC-PBP (Thermo Scientific PV4406), 100nM calmodulin, 10 nM calcineurin, and CLMB peptide at the indicated concentrations (Millipore Sigma 208694-1MG)). Reactions were initiated by adding Ca^2+^ and CaM to their final concentrations. Reactions took place in a total of 50 µL in a corning black 96 well vessel and were monitored continuously via fluorescence emitted by phosphate bound MDCC-PBP (EX:400/ EM:460). Phosphate standards were used to convert fluorescence intensity to phosphate concentration.

### Kinetic Modeling

Dephosphorylation data were fit by simulation and non-linear regression using Kintek Explorer™. Sigma values for each individual replicate were obtained using the double exponential afit function. All replicates were simultaneously fit to the models described in Supplemental Fig.s 8 (a,b). Curve offsets were capped at a maximum offset of 50nM. Substrate concentration was allowed to vary by 15% during fitting. Initial parameter values were chosen based on previously published rates, when available. The final offset and initial substrate concentration values for each curve, along with fitting information, are shown in Supplemental Tables 1-3. Final graphs display averaged (n ≥ 5) phosphate released at each timepoint with the corresponding fit. Simulated data were exported using 200 simulation steps and plotted using GraphPad Prism.

### Statistical analysis

Statistical analysis was done using Graphpad Prism version 10.2.2. All data is displayed as representative images or mean values with standard deviation error bars unless otherwise noted. All data represent three independent experiments unless otherwise indicated in figure legends. Image analysis of CLMB-calcineurin colocalization (Fig. 2C) was conducted using the following number of cells: CLMB_WT_ = 525, CLMB_IxITMut_ = 641, CLMB_LxVMut_ = 500, CLMB_T44A_ = 517. Statistical significance was calculated using one-way analysis of variance (ANOVA) with corrections for multiple comparisons as indicated.

## Data availability

All original scans of immunoblots, immunofluorescence images, and AlphaFold predictions can be accessed through Mendeley Data (to be made available upon publication). Plasmids and reagents will be shared upon request. NMR chemical shift assignments are deposited in the Biological Magnetic Resonance Bank (BMRB) under accession code 52641.

## Acknowledgements

We thank Norman Davey and members of the Cyert and Arthanari labs, in particular Angela Barth and Sneha Roy, for helpful discussion. We are especially grateful to Rick Russell (UT Austin) for feedback on the kinetic modeling and thank Callie Wigington and Makena Pule for their early contributions to the project. We thank Ilya Bezprozvanny (UT Southwestern) for suggesting the name, calcimembrin.

## Author Contributions

MSC, DAB, HA and JCR conceived the studies. SQ and TV provided early NMR analysis of CLMB-calcineurin complexes. DAB, supervised by MSC, carried out experiments and analyzed data for Figures 1, 2, and 5, and prepared figures. RYM, supervised by MSC, carried out experiments and analyzed data for Supplementary figures 1 and 2a. JCR, supervised by HA, carried out experiments and analyzed data for Figures 3 and 4. TV carried out secondary structure analyses. DAB and MSC wrote the manuscript in consultation with HA and JCR.

