## Supplementary figures and tables for "A novel motif in calcimembrin/C16orf74 dictates multimeric dephosphorylation by calcineurin"

### SUPPLEMENTAL FIGURE LEGENDS

**Supplemental Fig. 1 S-acylation of $CLMB_{G2A}$ is detected in the presence of PalmB.** Representative immunoblot showing metabolic labeling of CLMB-FLAG (wild-type or G2A mutant) with palmitate analog (17-ODYA) in the presence or absence of PalmB, analyzed via CLICK chemistry with biotin-azide and detection with IRdye 800CW streptavidin (n = 3 independent experiments).

**Supplemental Fig. 2: CLMB is phosphorylated at T44 and contains a functional LxVP motif. a**. Electrophoretic mobilities of CLMB containing alanine substitutions in T44 ($CLMB_{T44A}$), T46 ($CLMB_{T46A}$) or both ($CLMB_{T44A,46A}$) shows that mutation of T44 but not T46 affects electrophoretic mobility. Thus, CLMB is predominantly phosphorylated at T44. **b.** CLMB peptide inhibits dephosphorylation of RII peptide by calcineurin *in vitro* demonstrating the presence of a functional LxVP motif. N=2 independent experiments with 2 technical replicates for each experiment.

**Supplemental Fig. 3. Full length CLMB protein and $CLMB_{WT}$ peptide bind similarly to calcineurin and binding affinity is constant over pH range = 6.9 to 8.0. a.** Fluorescence polarization (FP) saturation binding curves of FITC-$pCLMB_{WT}$ incubated with serial dilutions of CNA/B (0 to 12.5 µM). Experiments were carried out using buffers of pH 6.9 (20 mM Hepes), 7.4 (PBS) or 8.0 (50mM Hepes) to obtain the following dissociation constants: $K_D$ = 3.56 (2.93-4.33) for pH 6.9; $K_D$ = 3.83 (3.07-4.76) for pH 7.4; $K_D$ = 3.41 (2.40-4.82) for pH 8.0. **b.** Relative fluorescence units obtained for the different pH buffers used. FP data were fitted to a one-site binding model (mean ± s.d., n = 2 independent experiments). **c.** Competitive binding assay of unlabeled $CLMB_{WT}$ peptide and full length CLMB protein titrated against 5nM FITC-$pCLMB_{WT}$ peptide, 5 nM in the presence of 0.06 µM CNA/B. Data are mean ± s.d. for n = 2 independent experiments. **d.** Table displaying the experimentally obtained $IC_{50}$ values and converted $K_i$ values from **c**.

**Supplemental Fig. 4 CNA/B but not CNA engages the LxVP motif when CLMB is in excess.**
**a** CLMB amino acid sequence. Assigned residues are highlighted in green; non-assigned residues are highlighted in blue; prolines are represented in grey. **b** Predicted secondary structure for CLMB analyzed using TALOS-N (*1*). β-strand propensity plotted in green, α-helix propensity in red, and coil propensity in grey. **c.** Predicted secondary structure for CLMB analyzed using CheSPI (*2*). β-strand propensity plotted in green, and α-helix propensity in red. **d.** Expanded views of CLMB with a 2:1 stoichiometry calcineurin A/B and calcineurin A around selected assigned peaks comprising the LxV and IxIT motifs. Free CLMB spectra (red) are overlayed with the spectra of CLMB with CNA/B (blue) and CNA (black).**e** Expanded views of the glycine region of CLMB. (left) Spectra overlay of apo CLMB (red) with CNA/B (blue) in a 2:1 stoichiometry; (right) Spectra overlay of apo CLMB (red) with CNA (black) in a 2:1 stoichiometry.

**Supplemental Fig. 5 Blocking PxIxIT site highlights LxV-mediated binding of CLMB to CNA/B. a**. Free CLMB (50µM) spectra (red) overlayed with CLMB and $pCLMB_{LxVMu}$t (40µM) (blue). **b**. Expanded views of free CLMB (blue) and CLMB plus CNA/B (2:1) (red), with both conditions including $pCLMB_{LxVMut}$ (40µM). Areas around assigned peaks comprising the LxV and IxIT motifs, as well as CLMB N-terminus are shown. **c.** Expanded views of free CLMB (blue) and CLMB plus CNA/B (1:1) (red), with both conditions including $pCLMB_{LxVMut}$ (40µM). Areas around assigned peaks comprising the LxV and IxIT motifs, as well as CLMB N-terminus are shown.

**Supplemental Fig. 6 AlphaFold 3 predicts PxIxIT and LxVP binding at 2:1 CLMB to calcineurin. a** AlphaFold 3 prediction of Calcineurin (CNAtrunc/CNB) interacting with two copies of unmodified CLMB (left) or CLMB phosphorylated at Thr44 (right). CNAtrunc contains residues 1-391 of CNA and excludes the calmodulin-binding and regulatory regions. Surface representations of CNA (grey) and CNB (light blue) colored by chain with LxVP and PxIxIT binding surfaces and active site colored in white. CLMB cartoon representations and side chains are colored by pIDDT score with low confidence regions in red and high confidence in blue. **b** Expanded view of calcineurin-CLMB LxVP (left) and calcineurin-pCLMB (right), LxVP interactions and phosphorylated Thr44 (pT44) occupying the active site. **c** Expanded view of calcineurin-CLMB LxVP (left) and calcineurin-pCLMB (right) PxIxIT interactions.

**Supplemental Fig. 7 PxIxIT-binding but not catalytic activity is required for calcineurin to increase CLMB phosphorylation at T44 when overexpressed. a** Representative immunoblot of $CLMB_{WT}$-FLAG, but not $CLMB_{T44A}$-FLAG phosphorylation induced mobility shifts induced by CN overexpression. Performed using lysates inducibly expressing $CLMB_{WT}$-FLAG or $CLMB_{T44A}$-FLAG and transfected with empty vector (-) or CNA-GFP (+). Asterisks indicate phosphorylated CLMB. (n = 2 independent experiments) **b** Representative immunoblot of phosphorylation-based mobility shifts in $CLMB_{WT}$-FLAG induced by overexpression of $CNA_{WT}$-GFP and $CNA_{H151Q}$-GFP but not $CNA_{NIR}$-GFP. The $CNA_{H151Q}$ mutant is catalytically inactive (*3*). $CNA_{NIR}$ contains alanines substituting for PxIxIT-binding residues NIR (aa 330-332) (*4*). Performed using lysates of cells that inducibly express $CLMB_{WT}$-FLAG transfected with empty vector (control) or $CNA_{WT}$-GFP or the indicated CNA mutants. Asterisks indicate phosphorylated CLMB. (n = 3 independent experiments).

**Supplemental Fig. 8 A kinetic model for CLMB dephosphorylation by calcineurin. a** Kinetic model of $CLMB_{WT}$ dephosphorylation. E represents the activated calcineurin-CaM complex, S represents phosphorylated CLMB, P represents non-phosphorylated CLMB, and Q represents free phosphate. Numbers indicate the site at which CLMB has bound calcineurin, with 1 and 2 indicating PxIxIT and LxVP binding respectively. k1/k-1 (blue) describe phosphorylated PxIxIT binding. k2/k-2 (pink) describe LxVP binding. k3 (black) describes dephosphorylation. Non-phosphorylated PxIxIT binding was omitted from modeling as the associated rates were poorly constrained by the data. **b** Kinetic model of $CLMB_{IxITMut}$ dephosphorylation using the same definitions as in **a. c** Binding and catalytic constants with standard error as determined by Kintek explorer when simultaneously fitting $CLMB_{WT}$ and $CLMB_{IxITMut}$ to their respective models.

**Supplemental Fig. 9 Simulations reveal key relationships between rates that control CLMB dephosphorylation behavior. a**. Simulated data of $CLMB_{WT}$ and $CLMB_{IxITMut}$ dephosphorylation at the indicated peptide and calcineurin concentrations using the estimated rates shown in in Supplemental Fig. 8. **b**. Simulated data, as in a, with varied phospho-PxIxIT binding on and off-rates ($k_1$ and $k_{-1}$) but a constant ratio between the two ($K_D$). The rates range from 100-fold less to 100-fold more than the original values. The dashed box indicates plots that deviate significantly from the original fit (a). **c**. Simulated data, as with b, but instead varying the LxVP off-rate and $k_{cat}$ ($k_{-2}$ and $k_3$) while keeping a constant ratio between the two. **d**. Simulated data, as in a, in which the LxVP on and off-rates ($k_2$ and $k_{-2}$) and $k_{cat}$ ($k_3$) are varied but the second order rate constant (equation shown in figure) remains constant. For each plot, one value is increased 10-fold (bolded), another value is adjusted to compensate for that change (bold and underlined), and the final value is not changed. For **b-d**, the dashed box indicates plots that deviate significantly from the original fit (a) and thus do not agree well with experimental data.

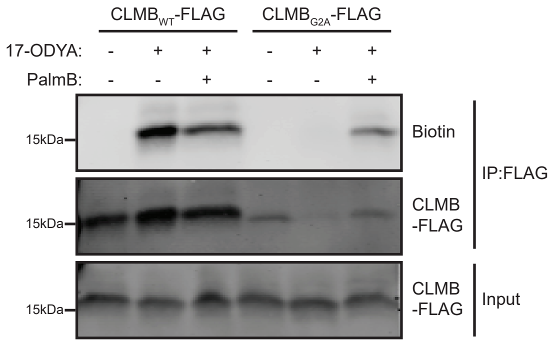
CLMB$_{WT}$-FLAG
CLMB$_{G2A}$-FLAG
17-ODYA: - + + - + +
PalmB: - - + - - +
15kDa
Biotin
15kDa
CLMB -FLAG
IP:FLAG
15kDa
CLMB -FLAG
Input

a

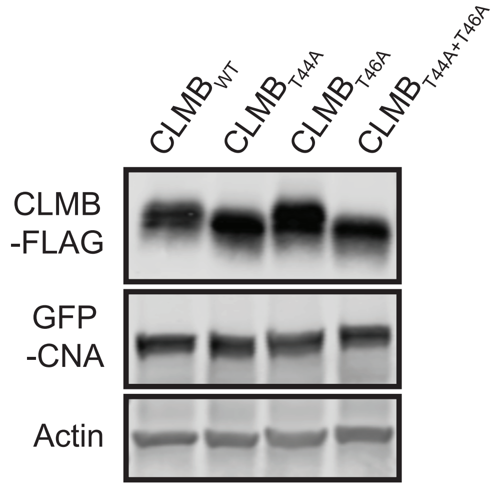

$CLMB_{WT}$
$CLMB_{T44A}$
$CLMB_{T46A}$
$CLMB_{T44A+T46A}$
CLMB
-FLAG
GFP
-CNA
Actin


b

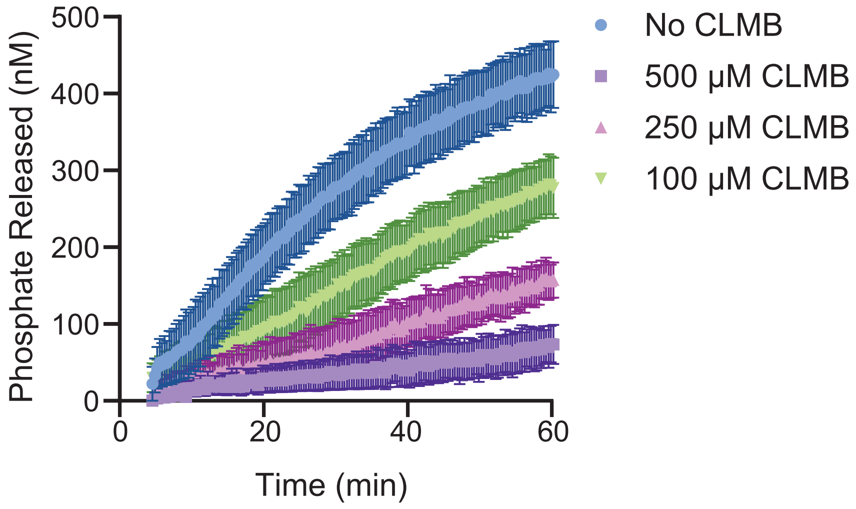

Phosphate Released (nM)
500
400
300
200
100
0
0
20
40
60
Time (min)
No CLMB
500 μM CLMB
250 μM CLMB
100 μM CLMB

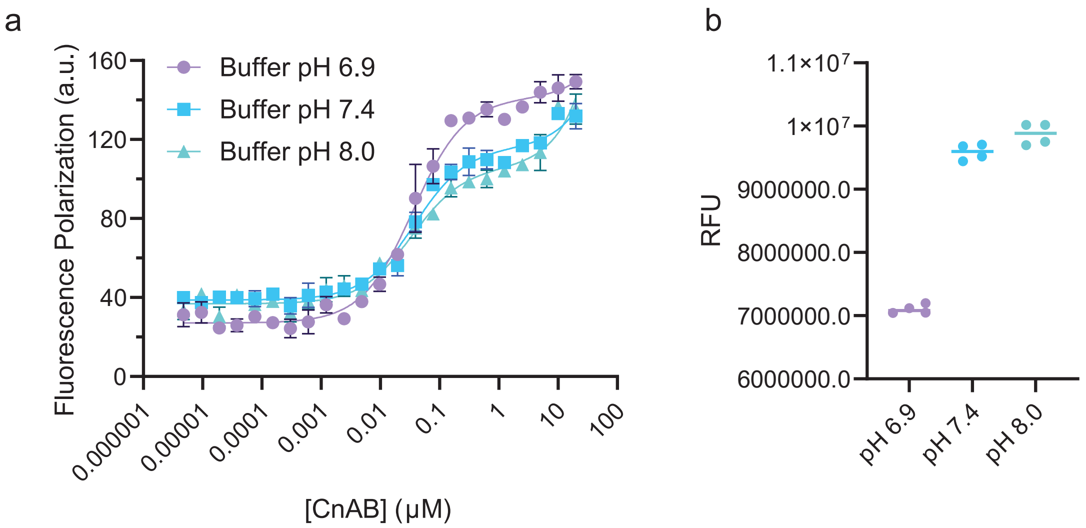


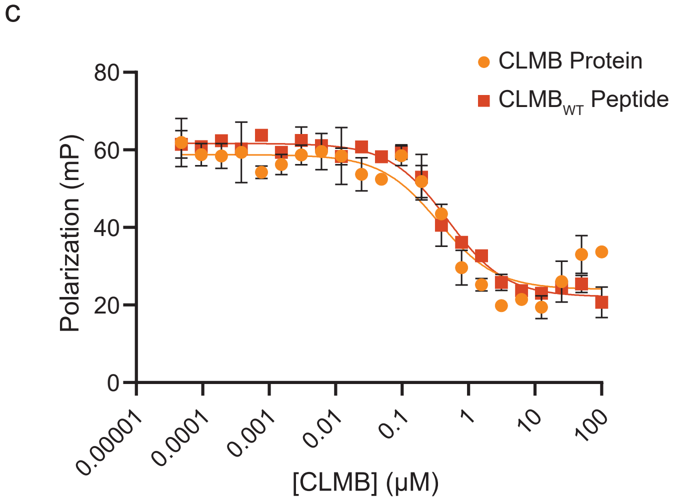


d

| Competitor | FITC-pCLMB$_{WT}$ | |
|---|---|---|
| | $IC_{50}$(nM) | $K_i$ (nM) |
| CLMB$_{WT}$ Peptide | 485.4 ± 61.3 | 476.7± 60.2 |
| CLMB Protein | 396.3 ± 109.1 | 389.2± 107.1 |

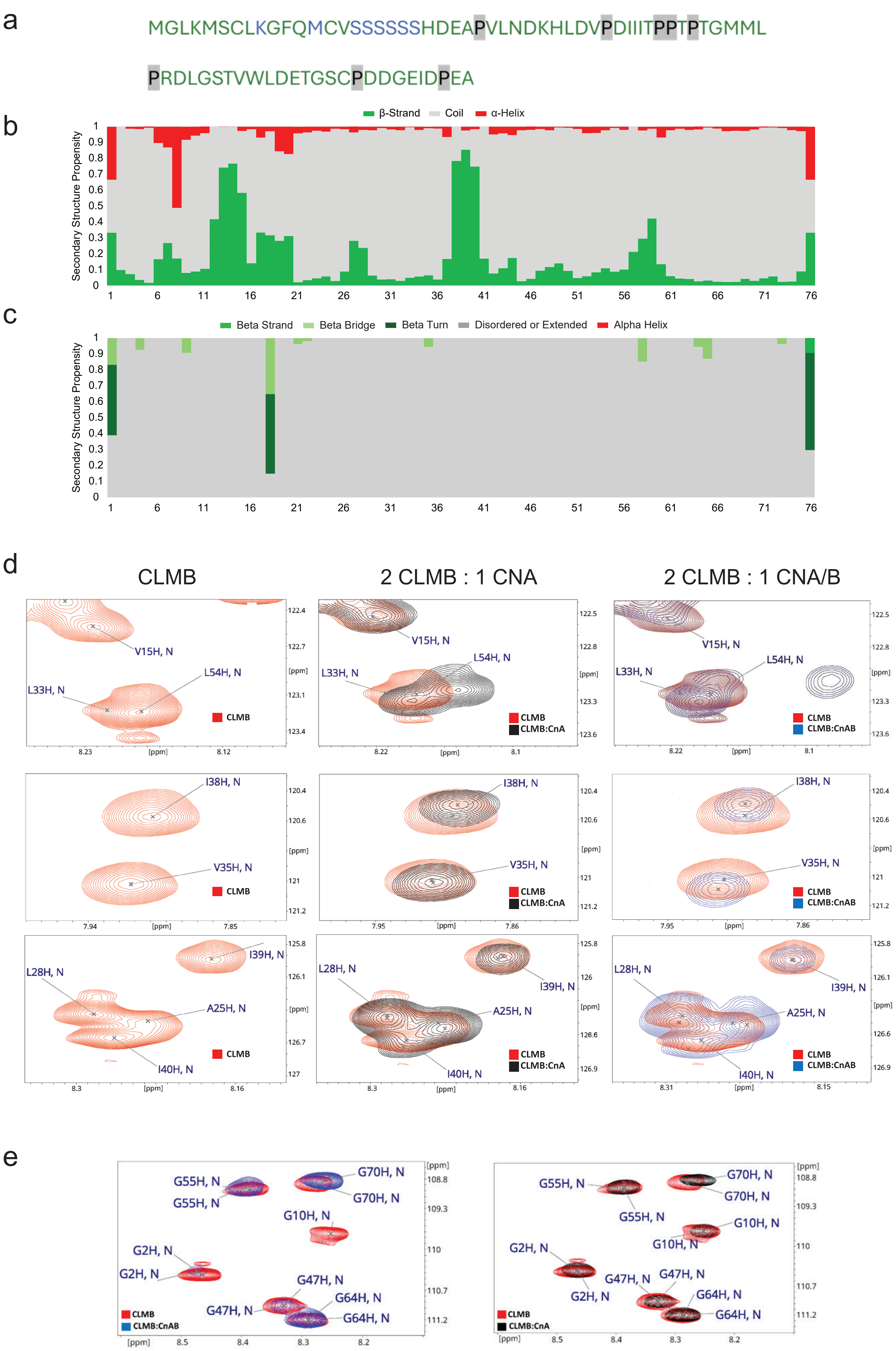

a
MGLKMSCLKGFQMCVSSSSSSHDEAPVLNDKHLDVPDIIITPPTPTGMML
PRDLGSTVWLDETGSCPDDGEIDPEA
b
β-Strand
Coil
α-Helix
Secondary Structure Propensity
c
Beta Strand
Beta Bridge
Beta Turn
Disordered or Extended
Alpha Helix
Secondary Structure Propensity
d
CLMB
2 CLMB : 1 CNA
2 CLMB : 1 CNA/B
V15H, N
L54H, N
L33H, N
I38H, N
V35H, N
L28H, N
I39H, N
A25H, N
I40H, N
CLMB
CLMB:CnA
CLMB:CnAB
[ppm]
e
G55H, N
G70H, N
G10H, N
G2H, N
G47H, N
G64H, N
CLMB
CLMB:CnAB
CLMB:CnA
[ppm]

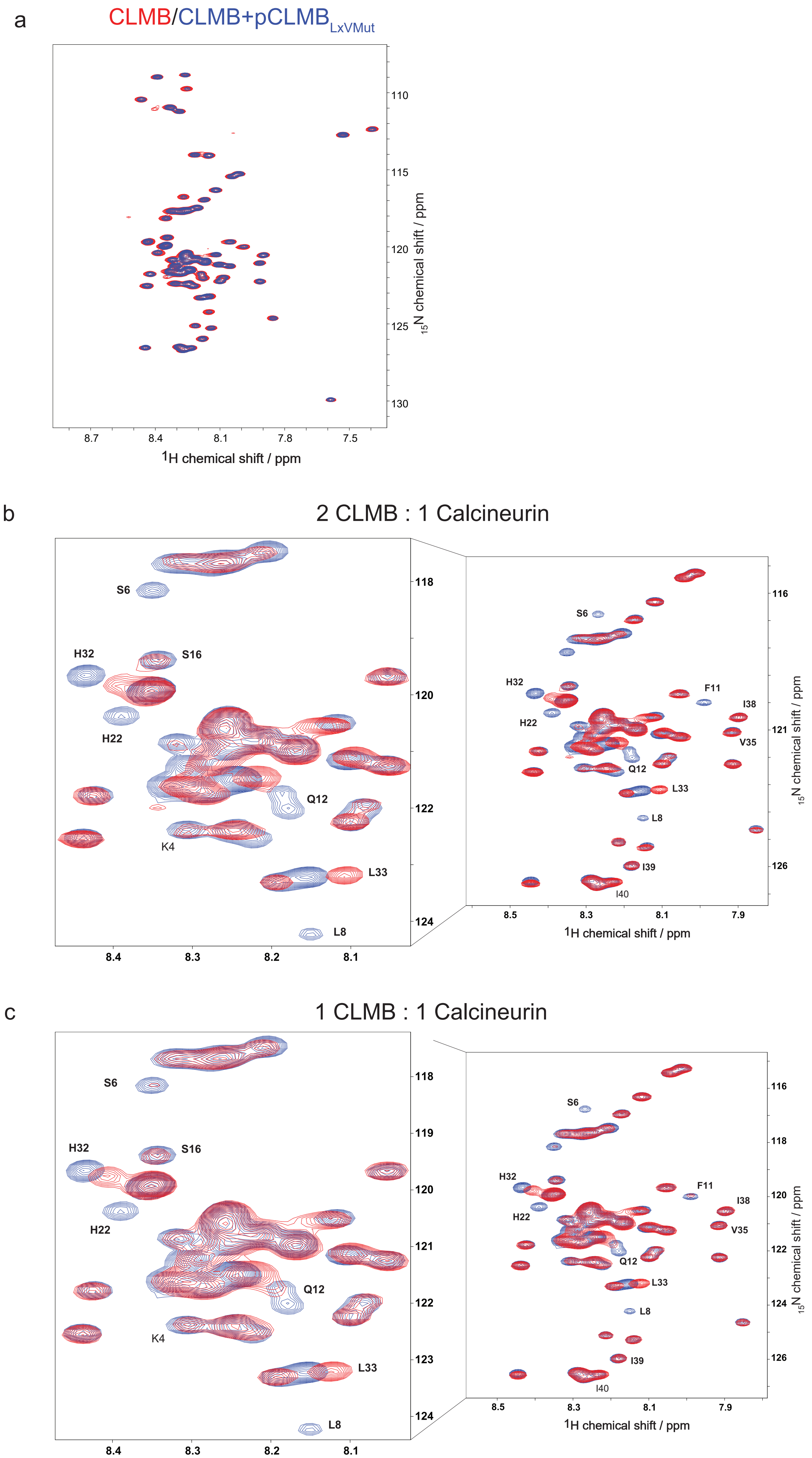
a
CLMB/CLMB+pCLMB LxVMut
1H chemical shift / ppm
15N chemical shift / ppm
b
2 CLMB : 1 Calcineurin
S6
H32
S16
H22
Q12
K4
L33
L8
F11
I38
V35
I39
I40
1H chemical shift / ppm
15N chemical shift / ppm
c
1 CLMB : 1 Calcineurin
S6
H32
S16
H22
Q12
K4
L33
L8
F11
I38
V35
I39
I40
1H chemical shift / ppm
15N chemical shift / ppm

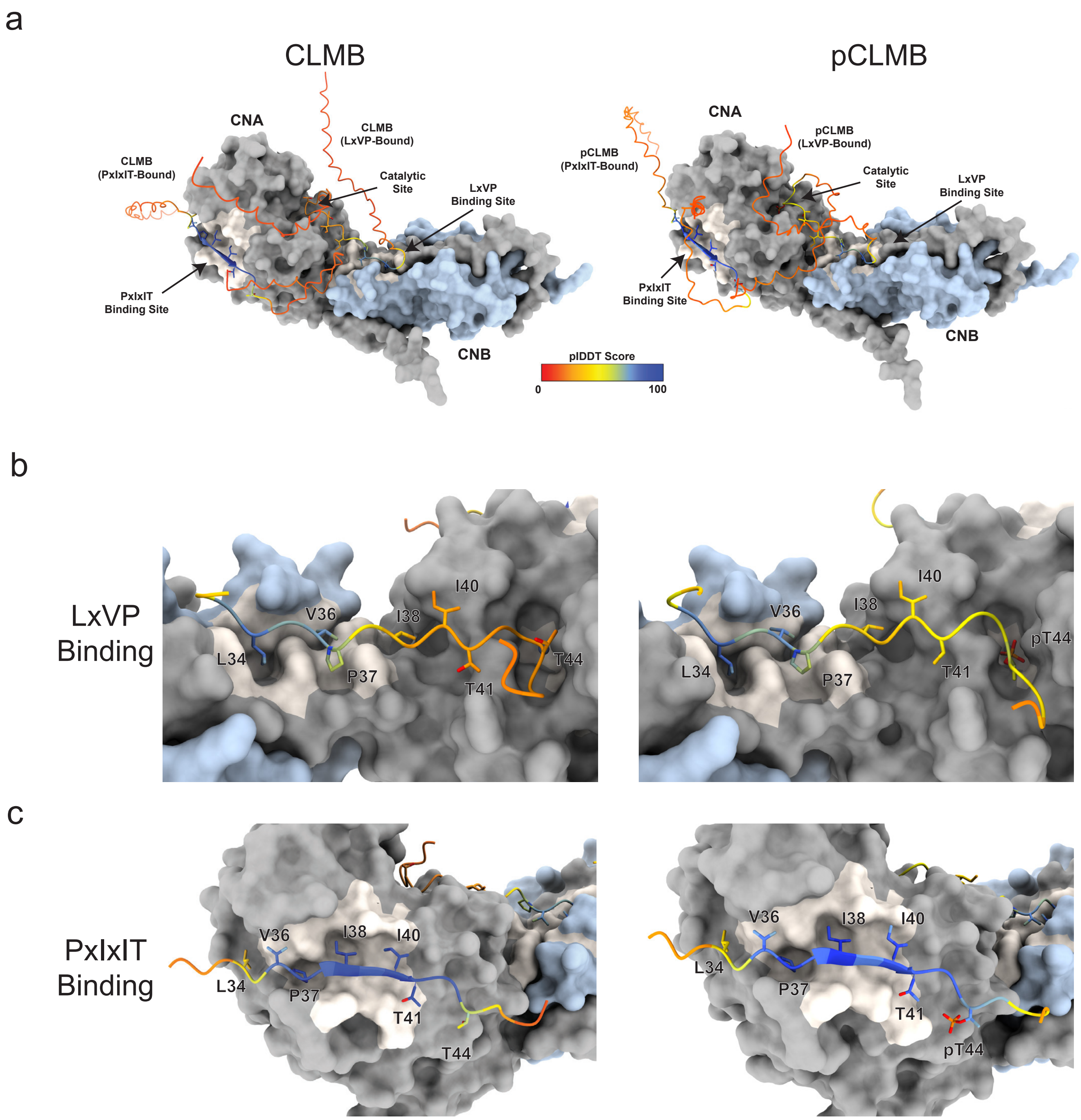
a
CLMB
pCLMB
CNA
CLMB (LxVP-Bound)
CLMB (PxIxIT-Bound)
Catalytic Site
LxVP Binding Site
PxIxIT Binding Site
CNB
pIDDT Score
0
100
pCLMB (LxVP-Bound)
pCLMB (PxIxIT-Bound)
b
LxVP Binding
L34
V36
P37
I38
I40
T41
T44
pT44
c
PxIxIT Binding

a

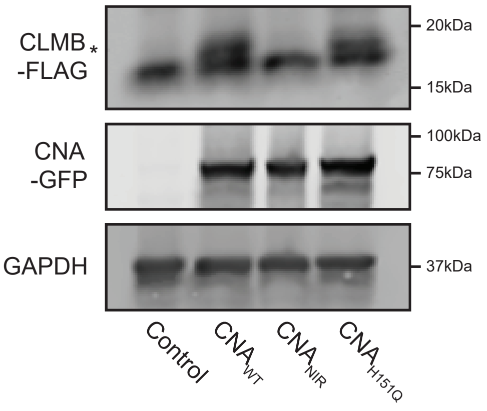
CLMB
-FLAG
*
20kDa
15kDa
CNA
-GFP
100kDa
75kDa
GAPDH
37kDa
Control
CNA WT
CNA NIR
CNA H151Q

a

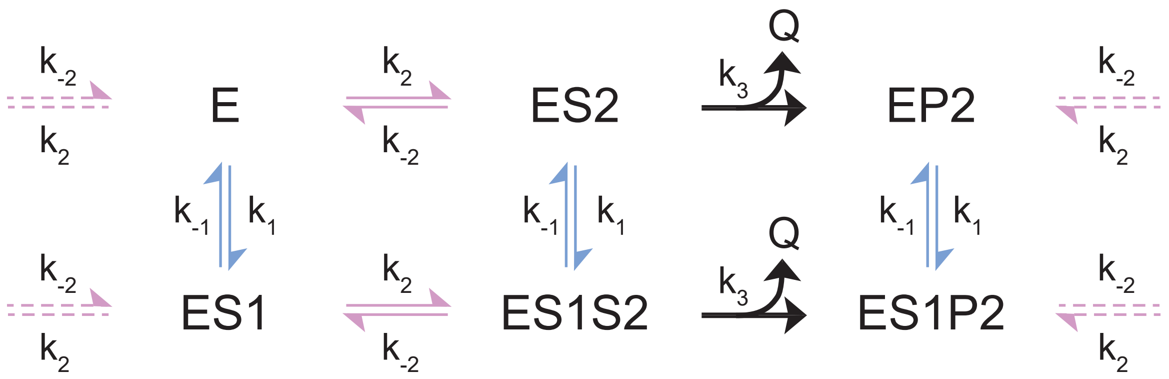


b

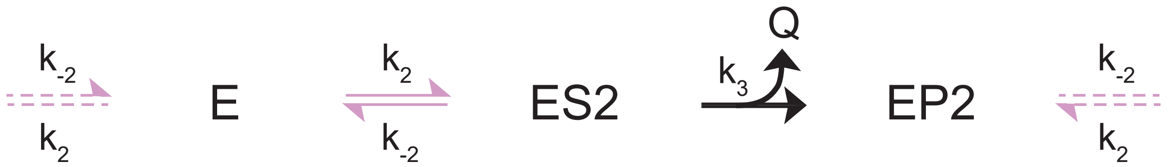


c

Binding Steps

| Rate | Binding Motif | $k_{on}$ ($M^{-1}$ $sec^{-1}$) | $k_{off}$ ($sec^{-1}$) |
|---|---|---|---|
| $k_1$ | Phospho-PxIxIT | $1.56 \pm 2.30 \times 10^{6}$ | $1.11 \pm 0.85 \times 10^{-2}$ |
| $k_2$ | LxVP | $4.86 \pm 4.65 \times 10^{4}$ | $8.44 \pm 3.94 \times 10^{-1}$ |

Catalytic Steps

| Rate | Reaction | $k_{cat}$ ($sec^{-1}$) |
|---|---|---|
| $k_3$ | Dephosphorylation | $6.69 \pm 2.85 \times 10^{-1}$ |

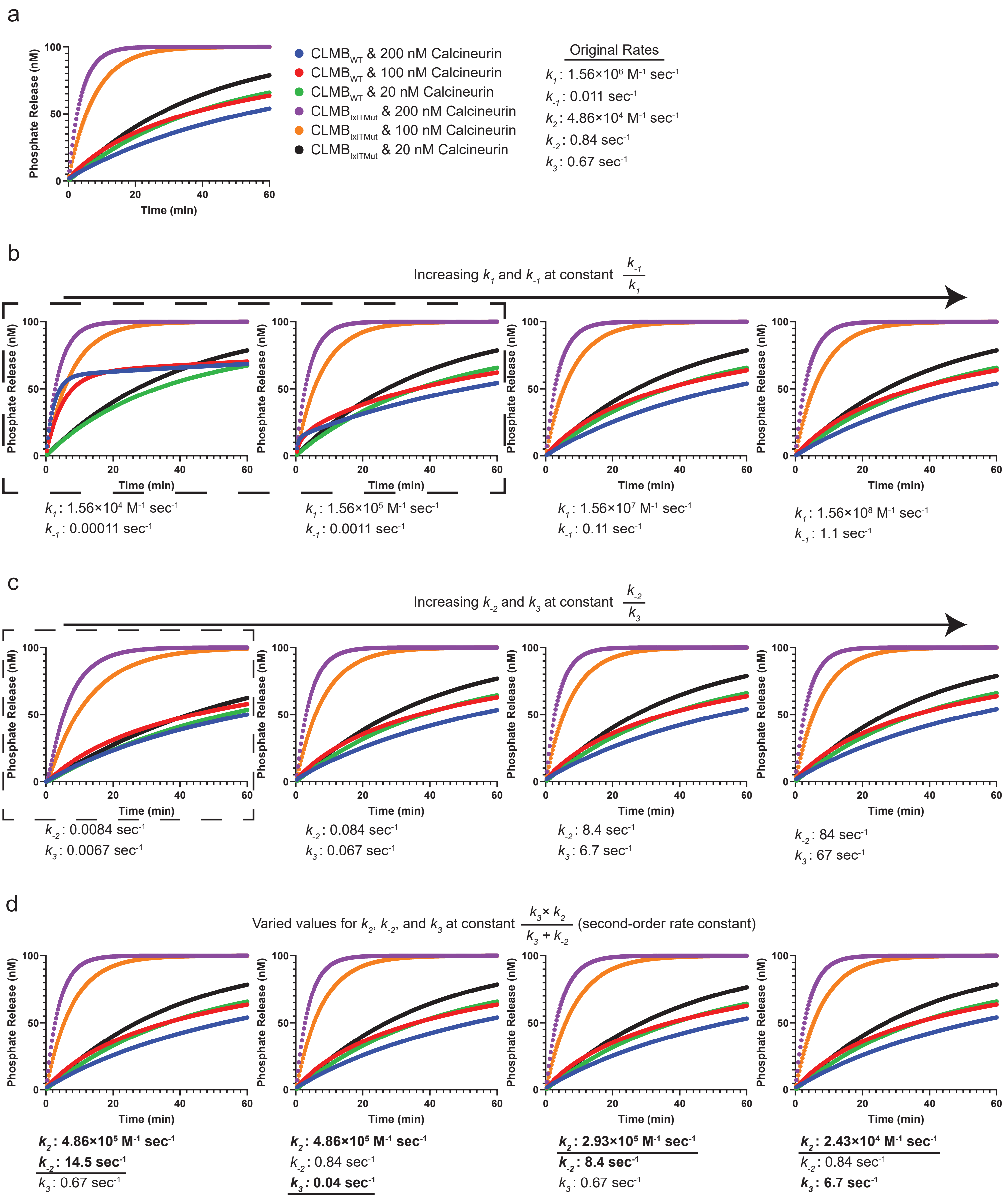
a
Phosphate Release (nM)
Time (min)
CLMB$_{WT}$ & 200 nM Calcineurin
CLMB$_{WT}$ & 100 nM Calcineurin
CLMB$_{WT}$ & 20 nM Calcineurin
CLMB$_{IxITMut}$ & 200 nM Calcineurin
CLMB$_{IxITMut}$ & 100 nM Calcineurin
CLMB$_{IxITMut}$ & 20 nM Calcineurin
Original Rates
$k_1$: 1.56×10$^6$ M$^{-1}$ sec$^{-1}$
$k_{-1}$: 0.011 sec$^{-1}$
$k_2$: 4.86×10$^4$ M$^{-1}$ sec$^{-1}$
$k_{-2}$: 0.84 sec$^{-1}$
$k_3$: 0.67 sec$^{-1}$
b
Increasing $k_1$ and $k_{-1}$ at constant $\frac{k_{-1}}{k_1}$
$k_1$: 1.56×10$^4$ M$^{-1}$ sec$^{-1}$
$k_{-1}$: 0.00011 sec$^{-1}$
$k_1$: 1.56×10$^5$ M$^{-1}$ sec$^{-1}$
$k_{-1}$: 0.0011 sec$^{-1}$
$k_1$: 1.56×10$^7$ M$^{-1}$ sec$^{-1}$
$k_{-1}$: 0.11 sec$^{-1}$
$k_1$: 1.56×10$^8$ M$^{-1}$ sec$^{-1}$
$k_{-1}$: 1.1 sec$^{-1}$
c
Increasing $k_{-2}$ and $k_3$ at constant $\frac{k_{-2}}{k_3}$
$k_{-2}$: 0.0084 sec$^{-1}$
$k_3$: 0.0067 sec$^{-1}$
$k_{-2}$: 0.084 sec$^{-1}$
$k_3$: 0.067 sec$^{-1}$
$k_{-2}$: 8.4 sec$^{-1}$
$k_3$: 6.7 sec$^{-1}$
$k_{-2}$: 84 sec$^{-1}$
$k_3$: 67 sec$^{-1}$
d
Varied values for $k_2$, $k_{-2}$, and $k_3$ at constant $\frac{k_3 \times k_2}{k_3 + k_{-2}}$ (second-order rate constant)
$k_2$: 4.86×10$^5$ M$^{-1}$ sec$^{-1}$
$k_{-2}$: 14.5 sec$^{-1}$
$k_3$: 0.67 sec$^{-1}$
$k_2$: 4.86×10$^5$ M$^{-1}$ sec$^{-1}$
$k_{-2}$: 0.84 sec$^{-1}$
$k_3$: 0.04 sec$^{-1}$
$k_2$: 2.93×10$^5$ M$^{-1}$ sec$^{-1}$
$k_{-2}$: 8.4 sec$^{-1}$
$k_3$: 0.67 sec$^{-1}$
$k_2$: 2.43×10$^4$ M$^{-1}$ sec$^{-1}$
$k_{-2}$: 0.84 sec$^{-1}$
$k_3$: 6.7 sec$^{-1}$

**Table 1: Kinetic model initial substrate concentration**

| Replicate | WT 200nM | WT 100nM | WT 20nM | IxIT 200nM | IxIT 100nM | IxIT 20nM |
|---|---|---|---|---|---|---|
| 1 | 97.6526 | 99.7556 | 99.863 | 107.574 | N/A | 97.851 |
| 2 | 100.62 | 99.9133 | 102.461 | 92.2289 | 101.199 | 99.3403 |
| 3 | 98.5251 | 96.3379 | 100.908 | 96.1978 | 96.4182 | 103.095 |
| 4 | 98.505 | 95.775 | 108.349 | 94.717 | 90.8177 | 104.816 |
| 5 | 101.198 | 104.424 | 105.746 | 100.597 | 104.029 | 109.348 |
| 6 | 99.6603 | 106.838 | 113.019 | 106.14 | 101.992 | 110.583 |

**Table 2: Kinetic model offset values.**

| Replicate | WT 200nM | WT 100nM | WT 20nM | IxIT 200nM | IxIT 100nM | IxIT 20nM |
|---|---|---|---|---|---|---|
| 1 | 30.08±3.8 | 27.13±2.76 | 24.58±2.57 | 34.79±4.79 | N/A | 18.47±2.74 |
| 2 | 49.41±3.51 | 28.64±2.63 | 24.08±2.53 | 0.36±4.17 | 16.97±3.45 | 18.64±2.69 |
| 3 | 18.13±3.36 | 9.28±2.48 | 10.83±2.46 | 2.2±4.07 | 0.32±3.23 | 13.39±2.73 |
| 4 | 30.42±3.32 | 5.62±2.47 | 27.04±2.59 | 1.32±3.98 | 0.13±3.12 | 13.44±2.75 |
| 5 | 22.87±4.21 | 33.59±3.06 | 8.9±2.8 | 16.54±5.18 | 27.55±3.97 | 22.29±3.09 |
| 6 | 24.72±4.13 | 44.56±3.11 | 49.42±2.92 | 26.98±5.12 | 26.17±3.9 | 24.5±3.1 |

**Table 3: Kinetic model fitting information**

| afit | double exponential |
|---|---|
| Sigma W.R.T. fit | 6.92531 |
| Chi2 | 756.41 |
| Chi2/DoF | 1.52455 |
| p-Value | 0 |
| Chi2 Thresh | 0.97336 |

**Table 4: Primers**

| | | |
|---|---|---|
| CLMB-GFP -> CLMB-FLAG | Forward Primer | 5'-TCACAAGGATCCGGCAGTGGCAGTGACTACAAAGACGATGACGATAAATAACTCGAGTTGTGA-3' |
| | Reverse Primer | 5'-TCACAACTCGAGTTATTTATCGTCATCGTCTTTGTAGTCACTGCCACTGCCGGATCCTTGTGA-3' |
| CLMB IxIT -> AxAA SDM | Forward Primer | 5'-GAACGACAAGCACCTGGACGTGCCCGACGCCATCGCCGCGCCCCCCACCCCCACGGGCATG-3' |
| | Reverse Primer | 5'-CATGCCCGTGGGGGTGGGGGGCGCGGCGATGGCGTCGGGCACGTCCAGGTGCTTGTCGTTC-3' |
| CLMB LxV -> AxA | Forward Primer | 5'-CGACAAGCACGCAGACGCACCCGACATCATCATC-3' |

| | | |
|---|---|---|
| | Reverse Primer | 5'-GATGATGATGTCGGGTGCGTCTGCGTGCTTGTCG-3' |
| CLMB T44A | Forward Primer | 5'-CGCCCCCCGCACCCACGG-3' |
| | Reverse Primer | 5'-CCGTGGGTGCGGGGGGCG-3' |
| CLMB C7S | Forward Primer | 5'-GCTTATGGGGCTTAAGATGTCCTCCCTGAAAGGCTTTCAAATGTGTG-3' |
| | Reverse Primer | 5'-CACACATTTGAAAGCCTTTCAGGGAGGACATCTTAAGCCCCATAAGC-3' |
| CLMB C7S -> C7,14S | Forward Primer | 5'-GAAAGGCTTTCAAATGAGTGTCAGCAGCAGCAGCAGCAGC-3' |
| | Reverse Primer | 5'-GCTGCTGCTGCTGCTGCTGACACTCATTTGAAAGCCTTTC-3' |
| CLMB G2A | Forward Primer | 5'-CCGGACTCTAGCGTTTAAACTTAAGCTTATGGCTCTTAAGATGTCCTGCCTGAAAGG-3' |
| | Reverse Primer | 5'-CCTTTCAGGCAGGACATCTTAAGAGCCATAAGCTTAAGTTTAAACGCTAGAGTCCGG-3' |
| CLMB-FLAG -> CLMB-V5 | Forward Primer | 5'-TGAGAAAAGCTTATGGGGCTTAAGATGTCCTGCCTGAAAGGCTTTCAAATGTGTG-3' |
| | Reverse Primer | 5'-TGAGAACTCGAGTTAGGTGCTGTCCAGGCCCAGCAGGGGG-3' |
| CNAH151Q-GFP | Forward Primer | 5'-CCCCAAAACACTGTTTTTACTTCGTGGAAATCAGGAATGTAGACATCTAACAGAGTATTTCAC-3' |
| | Reverse Primer | 5'-GTGAAATACTCTGTTAGATGTCTACATTCCTGATTTCCACGAAGTAAAAACAGTGTTTTGGGG-3' |

**Table 5: Plasmids**

| **Construct** | **Backbone** | **Plasmid Number** |
|---|---|---|
| CLMB$_{WT}$-FLAG | pcDNA5/FRT/TO | 2801 |
| CLMB$_{IxITMut}$-FLAG | pcDNA5/FRT/TO | 2862 |
| CLMB$_{LxVMut}$-FLAG | pcDNA5/FRT/TO | 2861 |
| CLMB$_{T44A}$-FLAG | pcDNA5/FRT/TO | 2860 |
| CLMB$_{LxVMut+T44A}$-FLAG | pcDNA3 | 2852 |
| CLMB$_{C7S}$-FLAG | pcDNA5/FRT/TO | 2863 |
| CLMB$_{C7,14S}$-FLAG | pcDNA5/FRT/TO | 2880 |

| | | |
|---|---|---|
| $CLMB_{G2A}$-FLAG | pcDNA5/FRT/TO | 2864 |
| $CLMB_{WT}$-V5 | pcDNA5/FRT/TO | 2918 |
| $CLMB_{IxITMut}$-V5 | pcDNA5/FRT/TO | 3125 |
| $CLMB_{LxVMut}$-V5 | pcDNA5/FRT/TO | 3126 |
| CNA-GFP | pcDNA5/FRT/TO | 2683 |
| $CNA_{NIR}$-GFP | pcDNA5/FRT/TO | 2794 |
| $CNA_{H151Q}$-GFP | pcDNA5/FRT/TO | 3166 |

Table 6: Antibodies

**Primary Antibodies**

| Antigen | Host | Dilution for WB | Dilution for IF | Supplier | Product Number |
|---|---|---|---|---|---|
| FLAG | Mouse | 1:2500 | NA | Sigma-Aldrich | F3165 |
| FLAG | Rabbit | 1:1000 | 1:100 | Sigma-Aldrich | F7425 |
| GFP | Rabbit | 1:1000 | NA | Thermofisher | A-11122 |
| GFP | Chicken | NA | 1:100 | Novus Biologicals | NB100-1614 |
| GAPDH | Mouse | 1:2000 | NA | Santa Cruz Biotechnology | sc-365062 |
| V5 | Mouse | 1:5000 | NA | Thermofisher | R960-25 |
| Calnexin | Rabbit | 1:2500 | NA | Enzo | ADI-SPA-865 |
| GM130 | Mouse | 1:3000 | NA | Cell Signaling Technology | 12480S |
| αTubulin | Rabbit | 1:2000 | NA | Sigma-Aldrich | T9026 |

**Secondary Antibodies**

| Reactivity | Host | Dye | Application | Dilution | Supplier | Product Number |
|---|---|---|---|---|---|---|
| Rabbit | Goat | cf555 | IF | 1:100 | Biotium | 20232 |
| Chicken | Goat | cf488a | IF | 1:100 | Biotium | 20020 |
| Mouse | Goat | cf647 | IF | 1:100 | Biotium | 20281 |
| Rabbit | Goat | 800CW | WB | 1:15000 | Licor | 926-32211 |
| Mouse | Goat | 680RD | WB | 1:15000 | Licor | 926-68070 |

**Labeled Streptavidin**

| Dye | Application | Dilution | Supplier | Product Number |
|---|---|---|---|---|
| 800CW | WB | 1:15000 | Licor | 926-32230 |
